# A lateral protrusion latticework connects neuroepithelial cells and is regulated during neurogenesis

**DOI:** 10.1101/2021.11.04.467255

**Authors:** Ioannis Kasioulis, Alwyn Dady, John James, Alan Prescott, Pamela A. Halley, Kate G. Storey

## Abstract

Dynamic contacts between cells within the developing neuroepithelium are poorly understood but play important roles in cell and tissue morphology and cell signalling. Here, using live-cell imaging and electron microscopy we reveal multiple protrusive structures in neuroepithelial apical endfeet of the chick embryonic spinal cord, including sub-apical protrusions that extend laterally within the tissue, and observe similar structures in human neuroepithelium. We characterise the dynamics, shape, and cytoskeleton of these lateral protrusions and distinguish these structures from cytonemes/filopodia and tunnelling nanotubes. We demonstrate that lateral protrusions form a latticework of membrane contacts between non-adjacent cells, depend on actin but not microtubule dynamics and provide a lamellipodial-like platform for further extending fine actin-dependent filipodia. We find that lateral protrusions depend on the actin-binding protein WAVE1: mutant-WAVE1 misexpression attenuated protrusion and generated a round-ended apical endfoot morphology. However, this did not alter apico-basal cell polarity nor reduce tissue integrity. During normal neuronal delamination sub-apical protrusions were withdrawn, but mutant-WAVE1-induced precocious protrusion loss was insufficient to trigger neurogenesis. This study uncovers a new form of cell-cell contact within the developing neuroepithelium regulation of which prefigures neuronal delamination.

## Introduction

The patterning and production of differentiating cells within a proliferative epithelium is a central activity in tissue homeostasis and embryonic development. Recent advances have identified roles for cellular protrusions in these processes in a range of tissues including epithelia (Gerdes and Pepperkok, 2013; González-Méndez et al., 2019; Mattes and Scholpp, 2018). It is now clear that there are many types of cellular protrusion and that these vary in dimension, dynamics and involvement of specific cytoskeletal components. Such protrusions can also be distinguished by whether their protrusive behaviour results in cell-to-cell membrane contact, as exhibited by cytonemes/filopodia (Ramírez-Weber and Kornberg, 1999) or connection and the sharing of cytoplasmic content, including small organelles, characteristic of tunnelling nanotubes (Rustom et al., 2004). Cytonemes are long, fine, actin-dependent filopodia which can contact cells up to several hundred microns away and were first shown to mediate signalling at a distance in *Drosophila* (Cohen et al., 2010; De Joussineau et al., 2003; Hsiung et al., 2005; Rajan et al., 2009). Since then, a plethora of similar cellular protrusions, some containing microtubules at their base, have been described in other embryos and their functional significance is beginning to be elucidated (González-Méndez et al., 2019). In the developing vertebrate nervous system, this has been most closely addressed in zebrafish. Here fine actin-dependent filopodia deliver Wnt ligand to pattern the neural plate (Luz et al., 2014; Mattes et al., 2018; Stanganello et al., 2015). Later, in the neural tube, the Notch ligand Delta, is presented by transient basally originating cell protrusions produced by newborn neurons to regulate the spatial and temporal production for spinal cord neurons (Hadjivasiliou et al., 2019). A correlation between filopodia and Notch ligand presentation has also been identified in the mouse cortex (Moore et al., 2020; Nelson et al., 2013), while other work supports a dependence on apical adherens junction localised Notch ligands in chick and mouse neuroepithelium (Hatakeyama et al., 2014). The former findings with respect to Wnt signalling and others reporting presentation of further ligands or their receptors, including Sonic hedgehog, Bone morphogenetic protein and Fibroblast growth factor, in non-neural contexts (Huang et al., 2019; Roy et al., 2014; Sanders et al., 2013) raise an alternative or complementary hypothesis to the notion that signal diffusion through extracellular space underlies establishment of signalling gradients which pattern developing tissues (Briscoe and Small, 2015; Kornberg, 2014; Wolpert, 2016). Whether similar cell protrusions exist and mediate patterning and neuronal differentiation or confer mechanical properties in the developing neuroepithelium in other model vertebrates or in human tissue has yet to be determined.

In the adult nervous system, cellular protrusions have been proposed to mediate pathogen spread in the form of prions and to facilitate progression of neurodegenerative diseases such as Alzheimer’s (Victoria and Zurzolo, 2017). A further related possibility is that cell protrusions propagate infectious agents within the developing neuroepithelium. These considerations highlight the importance of elucidating protrusive cell behaviour particularly at the apical surface of the neuroepithelium which is exposed to the fluid filled ventricle of the neural tube. The apical surface is made by the apical endfeet of neuroepithelial cells, which are linked together by N-Cadherin containing adherens junctions, essential for cell adhesion and tissue integrity (Hatta and Takeichi, 1986; Kadowaki et al., 2007; Meng and Takeichi, 2009). The pseudostratified nature of the neuroepithelium is characterised by interkinetic nuclear migration as neural progenitors progress through the cell cycle and divide at the apical surface (Miyata, 2008; Sauer, 1935). Within this proliferating cell population individual cells are then selected for neuronal differentiation by rising proneural gene expression and lateral inhibition mediated by Notch-Delta signalling (Henrique et al., 1995; Henrique and Schweisguth, 2019). Importantly, the transcription factor cascade that drives neuronal differentiation regulates the expression of *Cadherins* (Itoh et al., 2013; Rousso et al., 2012), reduction of which impacts adherens junctions and the linked intra-cellular actin network in the apical endfoot. These steps prefigure the delamination of newborn neurons from the ventricular surface in a process that involves local actin and microtubule cytoskeletal re-configuration (Camargo Ortega et al., 2019; Kasioulis and Storey, 2018; Kawaguchi, 2020) and in the chick and mouse neural tube, abscission of the apical membrane and dismantling of primary cilium (Das and Storey, 2014; Kasioulis et al., 2017), reviewed in (Kasioulis and Storey, 2018; Paridaen and Huttner, 2014). The presence of apical microvilli has been documented using electron microscopy (Shepard et al., 1997, 1993; Smith et al., 1982; Waterman, 1976) and immunofluorescence (Marzesco et al., 2005; Weigmann et al., 1997). Recent analyses have also identified sub-apical protrusions implicated in tissue elasticity underlying interkinetic nuclear migration (Shinoda et al., 2018). However, detailed characterization of the range of protrusive structures at the apical surface and extending sub-apically within the vertebrate neuroepithelium using live imaging approaches has yet to be undertaken.

Here we take advantage of the ability to transfect a mosaic of neuroepithelial cells in the chicken embryonic spinal cord to live image and monitor apical and sub-apical protrusion dimensions and dynamics. We further use electron microscopy to document apical endfoot protrusions in chicken and human neuroepithelium. Focusing on lamellipodia-like sub-apical lateral protrusions we parametrize these structures and demonstrate that they extend over several endfoot diameters and meet protrusions of non-neighbouring cells. Moreover, sub-apical lateral protrusions provided a platform for extension of fine filopodia, which we distinguish from cytonemes and tunnelling nanotubes. Distinct configurations of actin networks are regulated by distinct actin nucleating proteins of the WAVE family (Campellone and Welch, 2010; Sweeney et al., 2015; Yamazaki et al., 2005) and we show here that sub-apical lateral protrusions depend on WAVE1. We investigate the functional consequences of WAVE1 manipulation and identify a role for the regulation of the WAVE1-mediated actin network in neuronal delamination.

## Results

### Multiple protrusive structures emerge from neuroepithelial apical endfeet

To investigate the ultra-structural of neuroepithelial apical endfeet we used transmission electron microscopy (TEM), generating thin transverse sections of spinal cord (interlimb region) from chicken (HH stage 12) (Figures 1 A-C) and human (CS12) embryos (Figure 1D). This revealed key features in serial sections of the chicken tissue, including lumen protruding primary cilia (asterisks) and microvilli with a range of morphologies (blue arrowheads) (Figures 1 A-C). Focussing on contacts between neuroepithelial cells we identified thin plasma membrane that spread between adjacent endfeet (black arrow heads Figure 1C) apical to adherens junctions (white arrowheads), and sub-apical membrane protrusions (yellow arrowheads) that extend laterally over neighbouring cells below adherens junctions. These key sub-cellular structures could also be identified in human tissue (Figure 1D), which was distinguished by the presence of extensive microvilli, sub-apical lateral protrusions and prominent mitochondria (pink arrowheads).

**Figure 1.**
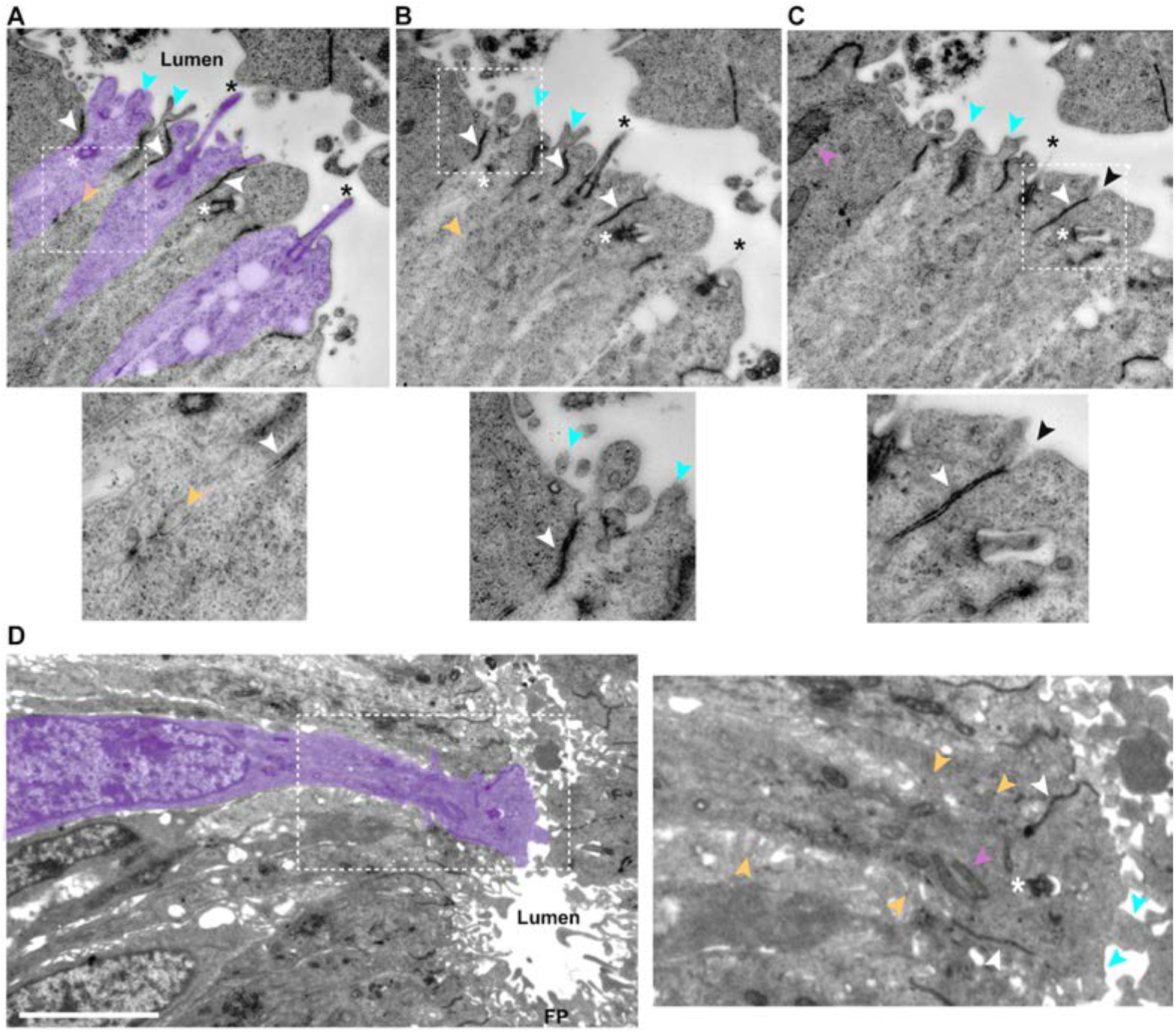
Ultra-structure of chicken and human embryo neuroepithelial apical endfeet. Serial transverse sections of chicken embryonic spinal cord (A-C), in A alternate endfeet are indicated in purple (excluding adherens junctions), white dashed line box is enlarged below each figure; (D)Transverse section though human embryonic spinal cord (CS12) in ventral region, including floorplate (FP), white dashed line box is enlarged in adjacent figure. Primary cilia tips in the lumen, black asterisks, and their basal bodies, white asterisks; adherens junctions (identified by electron dense material near abutting membranes), white arrowheads; microvilli, blue arrowheads; lateral protrusions, yellow arrowheads; mitochondria, pink arrowheads. Scale bar in D, 5 μm.

To investigate the dynamics of protrusive structures in apical endfeet, the neural tube of HH10-12 chick embryos was transfected with a plasmid encoding the membrane marker pm-eGFP and embryos cultured in ovo to stages HH17-18. This approach labelled cells in a mosaic fashion and allowed rapid live imaging of individual endfeet membrane dynamics from the apical surface (enface) and up to 5 μm basally into tissue explants.

A variety of apical membrane protrusions were observed, and these could be classified based on apico-basal position, size and dynamics. At the apical-most (lumen facing) surface short microvilli (aka micro-spikes) (0.66 ± 0.21 μm) (Shepard et al., 1997, 1993) consistent with those identified by TEM were most apparent, along with novel lamellipodia-like protrusive structures (1.44 ± 0.60 μm) which may correspond to the thin plasma membrane found between endfeet by TEM; microspikes and these apical lamellipodia-like protrusions (hereafter referred to as lamellopdia) were found in both live and fixed tissue (Figures 2A,B). Monitoring F-tractin localisation further confirmed that both the lamellipodia and the microvilli protrusions into the ventricle were actin rich (Movies 1-3). In addition, longer thin protrusions (5.34 ± 1.89 μm) that extended into the lumen and across neighbouring endfeet were observed at this apical-most surface. These filopodial-like protrusions were not apparent in our TEM analysis, but have been reported previously in SEM analyses (Shepard et al., 1997, 1993). Strikingly, these structures uniquely exhibited retrograde membrane movements (20-90 nm/s^-1^), consistent with the possibility of protein trafficking from the luminal space (Figures 2C-D, Movie 4).

**Figure 2.**
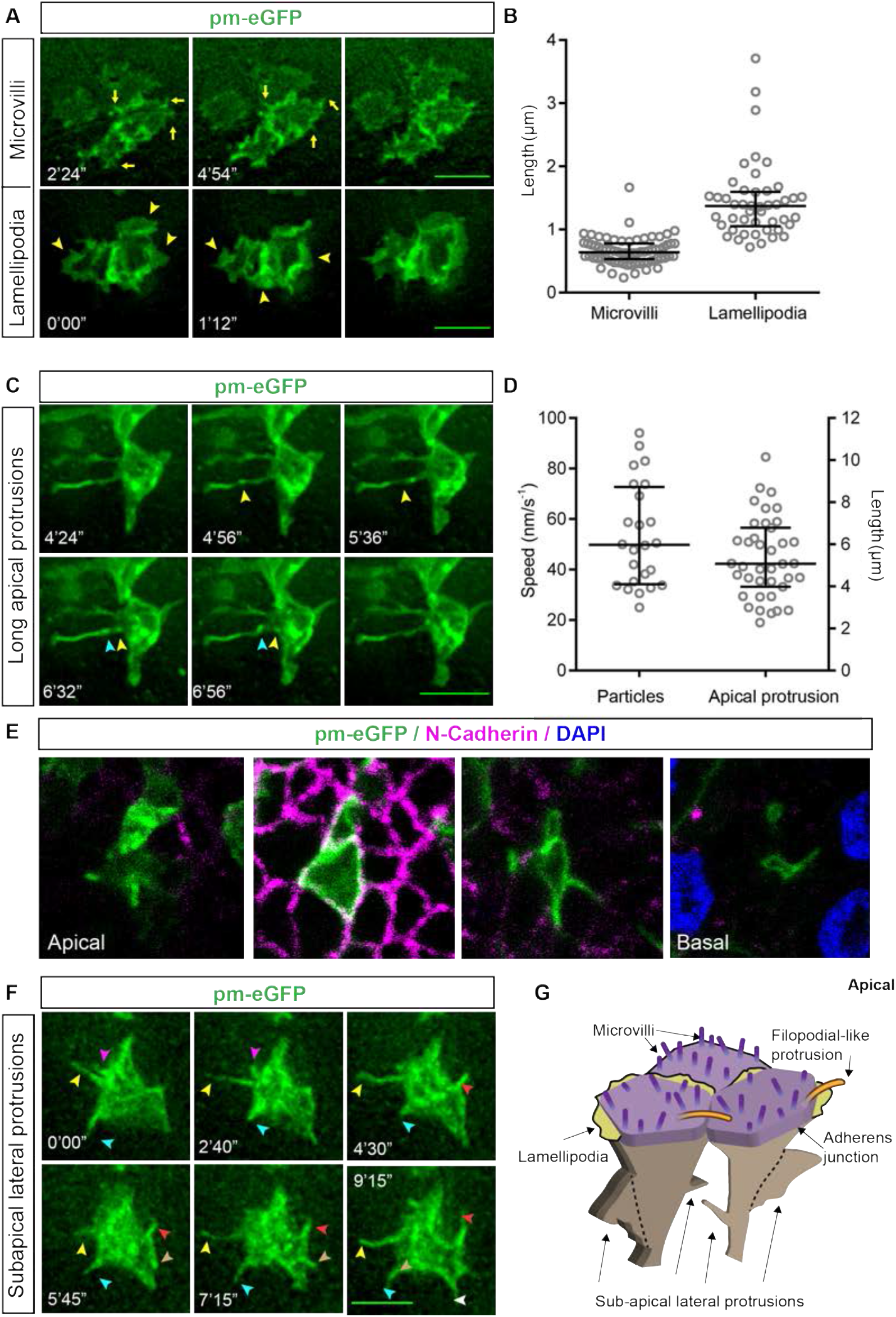
Live imaging analysis of apical and sub-apical protrusions extending from neuroepithelial apical endfeet. Enface live imaging of neuroepithelial cell apical end feet expressing the membrane marker pm-eGFP: (A) timepoints showing apical microvilli (yellow arrows) and lamellipodia (yellow arrowheads) at the apical surface; (B) quantification of microvilli and lamellipodial length (microvilli: 3 experiments, 7 explants, 28 cells, 70 microvilli; lamellipodia: 3 experiments, 7 explants, 46 cells); (C) long thin apical protrusions/ filopodia extending into the neural tube lumen, timepoints (and see movie 4); (D) quantification of surface protrusion length and speed of particle movement along the long thin apical protrusions (9 experiments, 14 explants, 39 cells); (E)Serial z stack images (apical to basal = right to left), showing sub-apical lateral protrusions emerging below N-cadherin expressing adherens junctions (pink); (F) timepoints showing sub-apical lateral protrusions, coloured arrowheads follow the dynamics of sub-apical protrusions over time; (G) Summary schematic of apical end foot protrusions, microvilli/microspikes, lamellipodia and long protrusions form at the apical surface (note primary cilia are not represented here). Sub-apical lateral protrusions emerge below the adherens junctions. Error bars in (B) and (D) are median with interquartile range. Scale bars in A, C, E, 5 μm.

Using this live imaging approach, a further set of membrane protrusions were observed, this time emerging sub-apically, beneath the N-cadherin expressing adherens junction (Figure 2E, Movie 5) and elongating lateral to the apico-basal axis, (Figure 2F, movie 6). These confirmed our EM observation of sub-apical lateral protrusions: these are similar to protrusive structures recently described in the developing mouse cortex (Shinoda et al 2018). In conclusion, the developing chick neuroepithelium contains at least three types of dynamic apical membrane protrusions, (microvilli, lamellipodia and filipodia), in addition to primary cilia, and exhibits sub-apical laterally extending protrusions, which hereafter we will refer to as lateral protrusions (Figure 2G).

### Characterisation of a dynamic lateral protrusions within developing neuroepithelium

To characterise lateral protrusions, we mis-expressed pm-eGFP alone or together with pm-mKate2 and live imaged these structures at short intervals to capture their dimensions (thickness and length) as well as speed and direction of movement.

Measurement of individual lateral protrusions revealed these to be thick (2-4 μm), short processes (most 2-3 μm, ranging up to 7 μm) (Figures 3A, B). To visualise lateral protrusion shape, explants fixed at the end of live imaging sessions were subject to higher magnification imaging using confocal microscopy to generate 3D models of apical end feet: these confirmed protrusion apico-basal position, thickness and length (Figures 3A, B and Movie 7). The direction of lateral protrusion movement was categorised as extension, retraction or lateral. Extension and retraction speeds ranged between 40-80 nm/s^-1^ with a few more rapid examples at 160-180 nm/s^-1^ (Figure 3C). In contrast, lateral movement was slower (20-60 nm/s^-1^) and detected most of the time (Figure 3D): this may reflect passive displacement influenced by shape change of neighbouring cells within the tissue.

**Figure 3.**
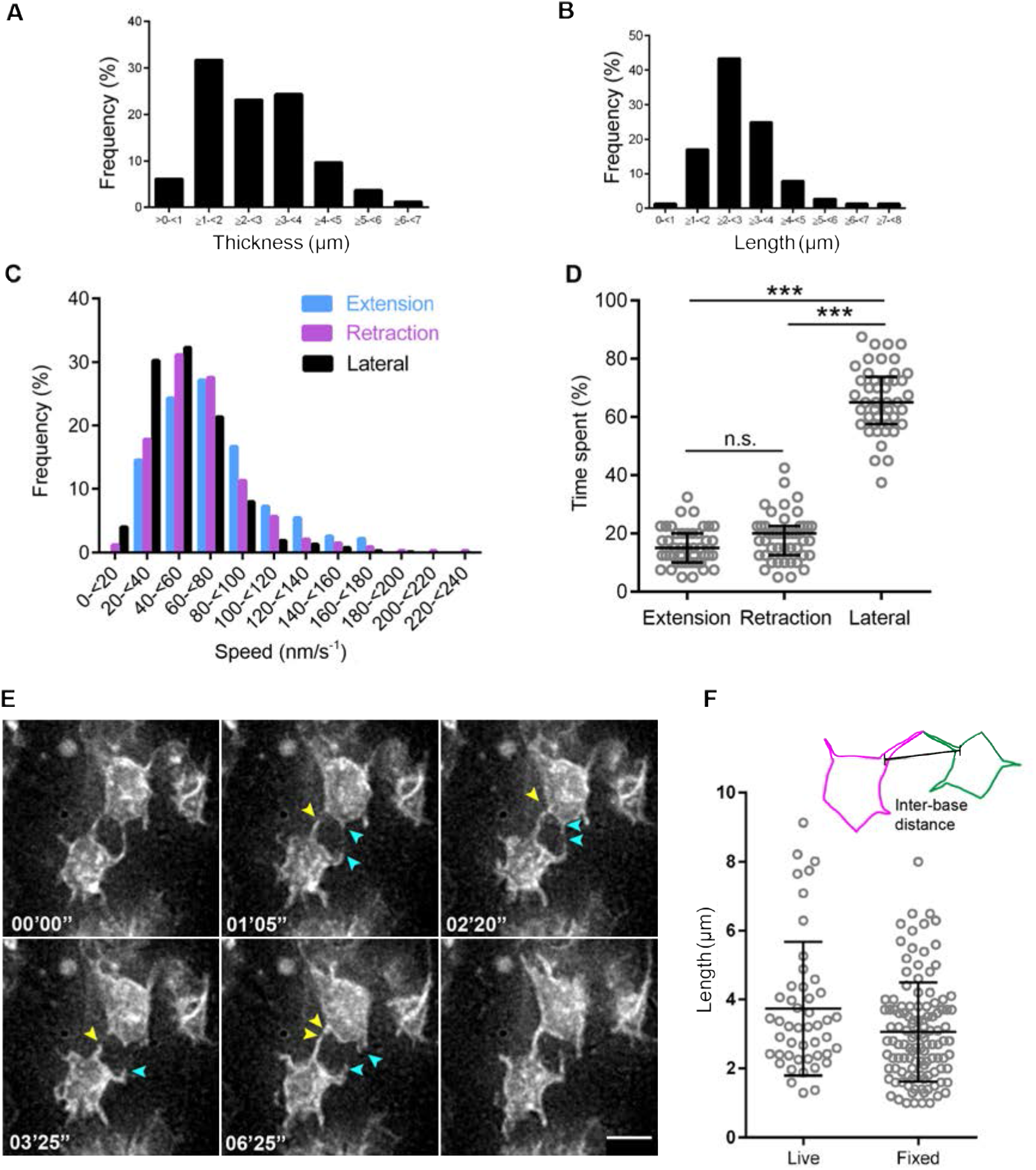
Characterisation of lateral protrusions. (A) Thickness of lateral protrusions and their frequency (3 experiments, 5 explants, 82 cells); (B) Protrusion length in contacting pairs (5 experiments, 9 explants, 48 pairs); (C) histogram of protrusion speed and frequency of extension, retraction, and lateral movement; (D) quantification of time individual protrusions extend, retract, or undergo lateral movement, total time analysed per protrusion is 10 minutes. One-way ANOVA test (Tukey’s multiple comparisons test), n.s.= not significant, p= 0.1786; *** p <0.0001); (E) timepoints from live imaging of lateral protrusions. Arrowheads follow the dynamics of protrusions over time and cells express the membrane marker pm-eGFP; (F) quantification of lateral protrusion distance from their base, measured in fixed and live-imaged tissue (Fixed: 3 experiments, 7 explants, 116 cells; Live: 5 experiments, 9 explants, 47 cells), lines represent mean with SD. Error bar in (B) is median with interquartile range, and (D) is mean with SD. Scale bar in (E) 5 μm.

A striking feature of lateral protrusions, made apparent by the mosaic transfection of cells, is their extension around immediate neighbouring endfeet and contact with similar protrusions from more distant cells within the tissue (Figure 3E, Movies 8 - 9). We measured the direct distance between the base of protrusions that contact each other and found that the majority were 1-5μm apart, while a subset reached across 5-10μm; similar findings were made in fixed tissue (live: 3.73 ± 0.28 μm, fixed: 3.1 ± 0.13 μm) (Figure 3F). As apical endfeet range from ~ 3-6 μm in diameter (Supplementary figure 1A) and lateral protrusions extend ~2-3 μm (Figure 3B), this indicates that most lateral protrusions make contact with protrusions from non-neighbouring cells. The extent of protrusion-protrusion contacts between non-neighbouring cells was next measured by analysing image Z-stacks. Most protrusion contacts ranged up to 2 μm (Supplementary Figure 1 B).

In addition, it was clear that lateral protrusion length varied from cell to cell in contacting pairs (Supplementary Figure 1C). Importantly, these protrusion-protrusion contacts were dynamic and could persist for more than 10 minutes (longest contact time recorded 22’55’’) (Figure 4A). To investigate contact dynamics in detail we transfected the neuroepithelium with a mixture of membrane markers pm-eGFP and pm-mKate2 (Figure 4B), generating a mosaic of cells expressing either one of these fluorophore-tagged proteins. Live imaging confirmed lateral protrusion extension around the curvature of neighbouring endfeet (Figure 4C and Movie 10) and revealed that protrusions from non-neighbouring cells meet tip to tip, extend along each other and also retract (Figures 4D – E, Movies 11 – 12). These behaviours suggest that lateral protrusions do not fuse but rather generate close membrane contacts between non-neighbouring cells.

**Figure 4.**
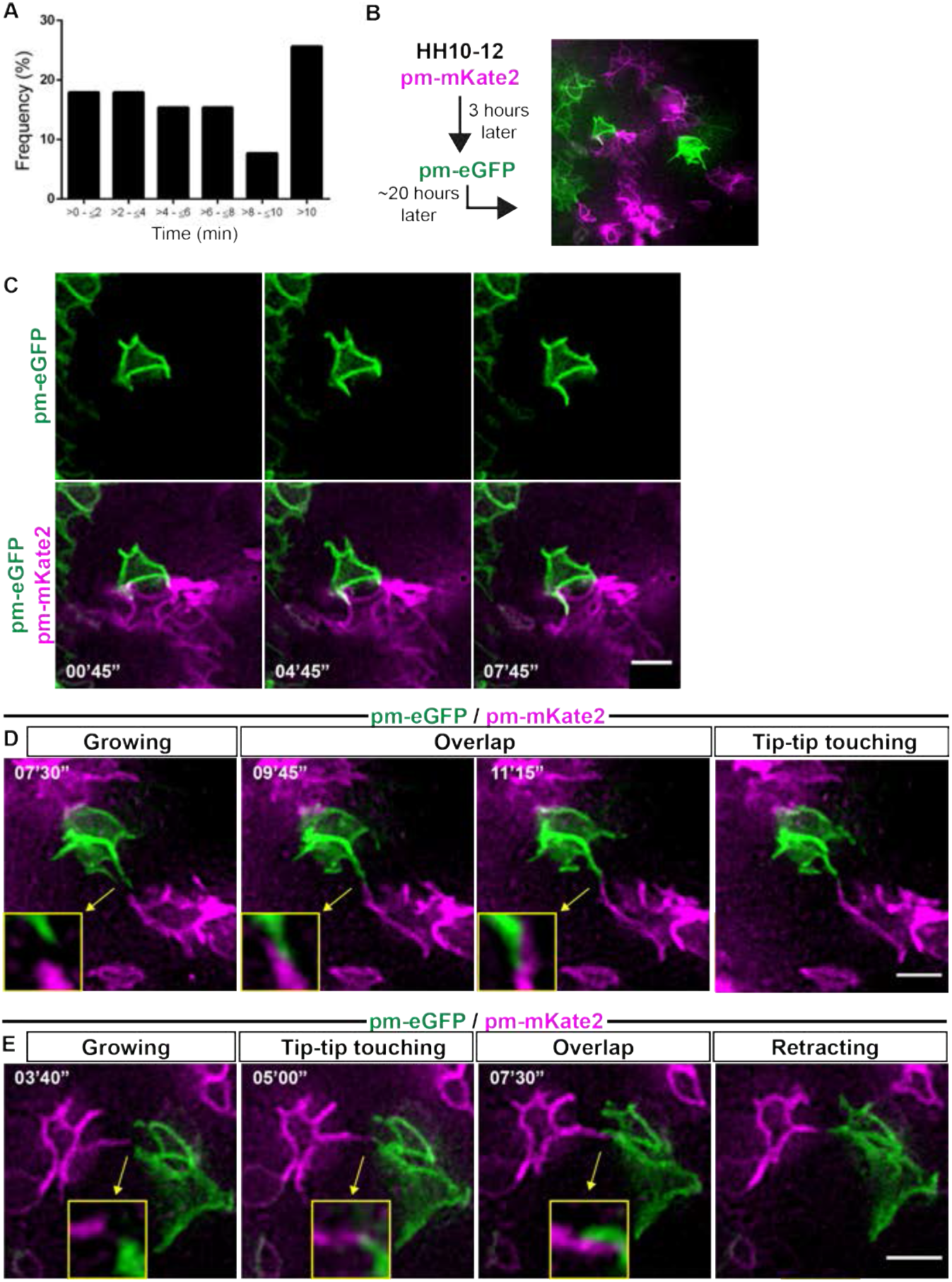
Lateral protrusion contacts from non-neighbouring cells extend along each other and can be long-lived. (A) Bar chart showing duration and frequency of protrusion contacts (5 experiments, 9 explants, 39 cell pairs); (B) strategy for generating mosaic pattern of pm-eGFP -pm-mKate2 pattern: cells are first transfected with pm-mKate2 at stage 10-12 and three hours later with pm-EGFP. Embryos are then left to develop to stage HH17-18 for live imaging; (C) timepoints showing that protrusions grow along the sides/edges of neighbouring cells; (D-E) timepoints showing protrusions from non-neighbouring cells contacting each other at their tips and extending along each other. Arrows and inset show the region magnified. Scale bars (C-E) 5 μm.

Together these observations identify a new form of cell-cell contact within the developing neuroepithelium mediated by a sub-apical latticework of laterally extending protrusions, which connect non-neighbouring cells within 1-2 endfoot diameters.

### Unique cytoskeletal architecture distinguishes lateral protrusions

To characterise the cytoskeletal architecture of lateral protrusions, plasmids expressing constructs that labelled actin (F-tractin-mKate2) or microtubule comets (EB3-sfGFP) were transfected along with membrane markers into the neural tube. F-tractin-mKate2 localisation followed the extension and retraction dynamics of the protrusions and was detected close to the protrusion tip (Figure 5A and Movie 13). EB3-sfGFP comets showed a unidirectional movement from base to tip in most protrusions (Figure 5B, Supplementary Figure 2A and Movie 14). Moreover, co-transfection of F-tractin-mKate2 and EB3-sfGFP showed that actin and microtubules elongate to the same extent within lateral protrusions (Supplementary Figure 2B-C).

**Figure 5.**
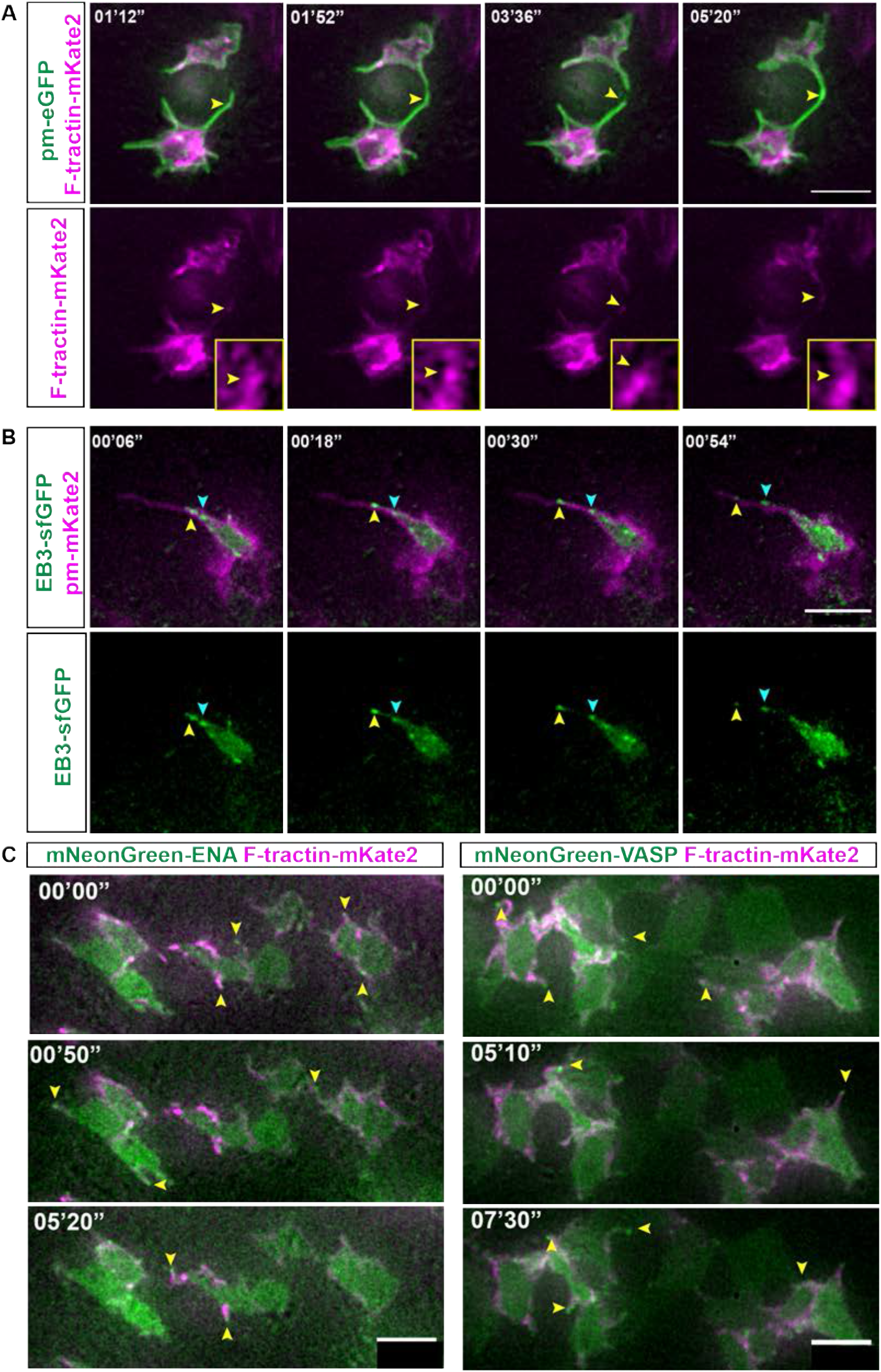
Actin, microtubules and ENA/VASP proteins localize to lateral protrusions. (A-C) Timepoints from live imaging sessions. (A) F-tractin-mKate2 follows the dynamics of membrane protrusions. Yellow arrowhead follows the F-tractin-mKate2 dynamics (2 experiments, 6 explants). Insets are of the region where the arrowhead is pointing. (B) Unidirectional microtubule movement from the base to the tip of the protrusion. Arrowheads follow the movement of EB3-sfGFP comets (2 experiments, 5 explants) (C) ENA and VASP localizing at the tip of protrusions (yellow arrowheads) (3 experiments, 10 explants). Scale bars (A-C) 5 μm.

Proteins involved in actin filament elongation in many different contexts include ENA and vasodilator stimulated phosphoprotein (VASP) (Dent et al., 2007; Gallop, 2020; Kwiatkowski et al., 2007; Lebrand et al., 2004). Ena/VASP are also proposed to support junctional actin assembly in tissue that is under frequent tension of myosin-induced contractility, including at the level of adherens junctions (Leerberg et al., 2014). To investigate if these proteins are localised to lateral protrusions, we mis-expressed (as above) mNeonGreen-ENA or mNeonGreen-VASP together with the actin marker F-tractin-mKate2 and both proteins were found at adherens junctions (Supplementary Figure 2D-E and Movies 15-16), and at lateral protrusion tips (Figure 5C, Movies 17 – 18). In contrast, a further tip localised regulator of filopodial elongation, unconventional Myosin X (MYO10), e.g. (Sanders et al., 2013), was detected uniformly in the cell cytoplasm but did not localise to lateral protrusion tips (Supplementary Figure 2F). However, following mis-expression Myosin X was found at the tip of a separate set of longer dynamics protrusions, the origin of which was difficult to determine (Movie 19). Along with their shorter and broader dimensions, these further findings distinguish lateral protrusions from elongating filopodia, which exhibit prominent tip localised Myosin X, and, consistent with their expression of Ena/VASP (Damiano-Guercio et al., 2020) indicate that lateral protrusions have lamellipodia-like characteristics.

To monitor lateral protrusion dynamics over a longer period we next increased the imaging interval, which reduces photobleaching and potential photo-toxic effects: images were captured once a minute for one hour. This regime revealed the emergence of thin filopodia from the lateral protrusions (Figure 6A and Movies 20 – 21). These lateral filopodia had a homogenous thickness and were extended on average 5 times per hour (Figure 6B-C). Lateral filopodia were longer than the lateral protrusions themselves and the combined length of protrusions and filopodia averaged 4.7μm (Figure 6D) and so together can extend within the neuroepithelium beyond their immediate cell neighbours. Together these findings show that both actin and microtubules extend within these lateral protrusions, that these structures have lamellipodia-like characteristics and that emerging filipodia extend the reach of lateral protrusions.

**Figure 6.**
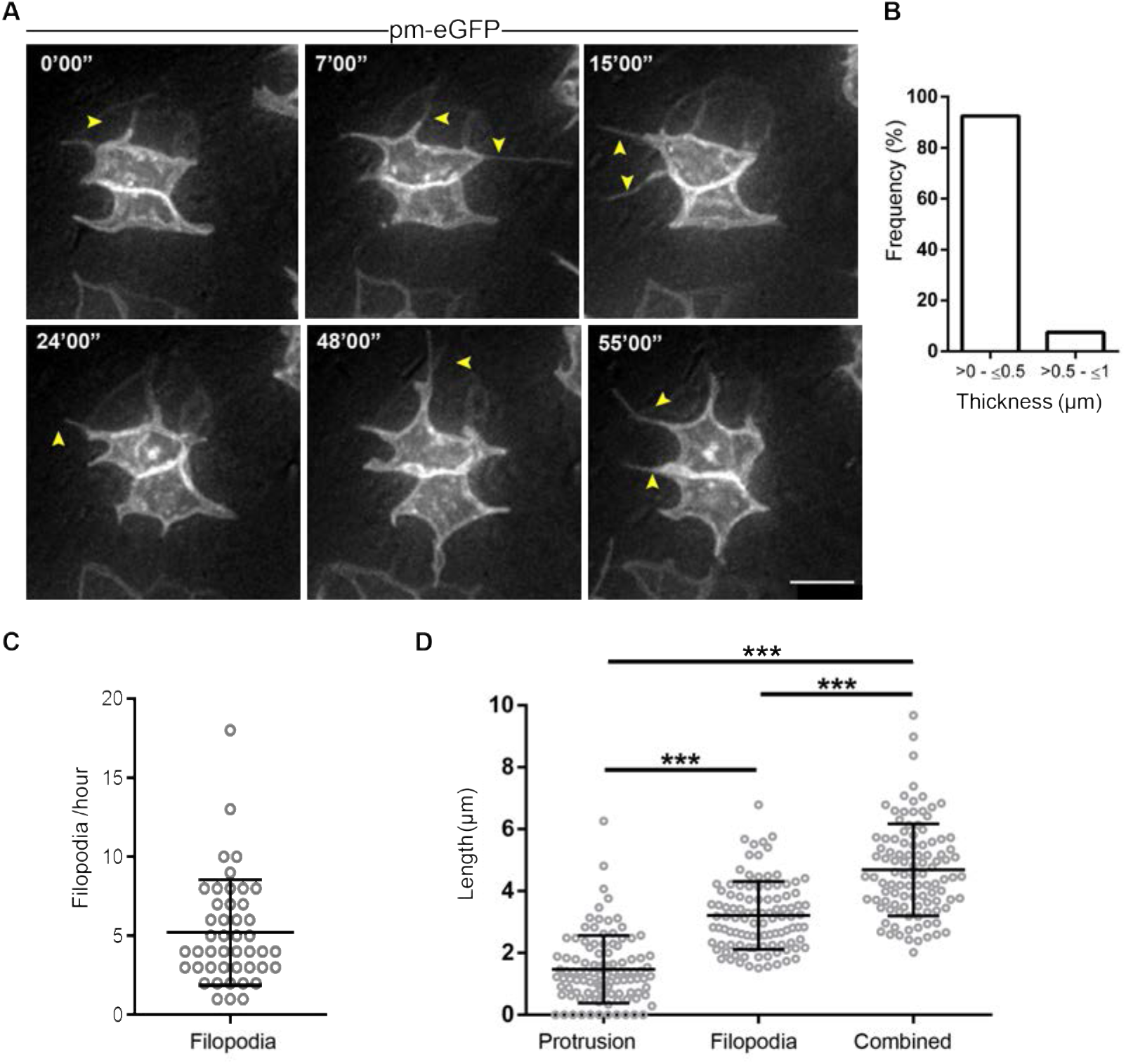
Filopodia emerge from lateral protrusions. (A) Live imaging timepoints showing presence of thin filopodial extensions (yellow arrowheads) emerging from lateral protrusions; (B) Bar chart showing filopodial protrusions are ~< 0.5μm in thickness (3 experiments, 5 explants, 65 filopodia); (C) Dot plot showing number of filopodia in one hour (5.2 ± 3.3 / hour) (4 experiments, 7 explants, 45 cells); (D) Filopodia are significantly longer than lateral protrusions, and combined filopodial-protrusion length extends lateral reach, (3 experiments, 5 explants, 27 cells) One-way ANOVA test (Tukey’s multiple comparisons test), ***p<0.0001. (C, D) Mean with standard deviation. Scale bar (A) 5 μm.

### Developing neuroepithelial cells do not share mitochondria via lateral protrusions

To investigate whether lateral protrusions act as tunnelling nanotubes within the neuroepithelium, we mis-expressed the mitochondrial marker mNeonGreen-TOMM20 and the membrane marker pm-mKate2. We then employed a fast imaging protocol to track both mitochondrial movements and cell membrane dynamics. Despite the dynamic nature of mitochondria within the cell cytoplasm, transfer between cells was never observed (Supplementary Figure 3A - B, Movie 22). We further generated and analysed electron microscopy images of the apical neuroepithelium of developing chick and human spinal cord for mitochondria localisation. These organelles were found in the apical endfoot in both species but were not located in lateral protrusions and appeared too large to enter these structures (Supplementary Figure 3 C-D). These live imaging and EM data, together with the finding that lateral protrusions from non-adjacent cells extend along each other (Figures 4D, E) suggest that the meeting of lateral protrusions does not lead to formation of cytoplasmic connection between cells.

### Lateral protrusions depend on actin but not microtubule dynamics and lateral filipodia are also actin dependent

To address whether actin or microtubule dynamics are necessary for the formation of lateral protrusions the membrane marker pm-eGFP was introduced into the neuroepithelium (as above) and explants were incubated in media containing Latrunculin-A (1μM), Taxol (10 μM) or control vehicle DMSO for one hour, followed by live imaging. Latrunculin-A binds to free actin dimers and depletes the pool of available actin for filament formation and maintenance, resulting in actin filament depolymerisation over time (Coué et al., 1987). Taxol binds bundles and stabilises microtubules, perturbing depolymerisation events (Schiff and Horwitz, 1980). Exposure to Latrunculin-A reduced lateral protrusion dynamics and lead to quantifiable inhibition of lateral filopodia formation, while Taxol had no effect on these structures (Figures 7A, B, movies 23 – 25). These data show that lateral protrusions and filopodia rely on actin but not microtubule dynamics for their extension.

**Figure 7.**
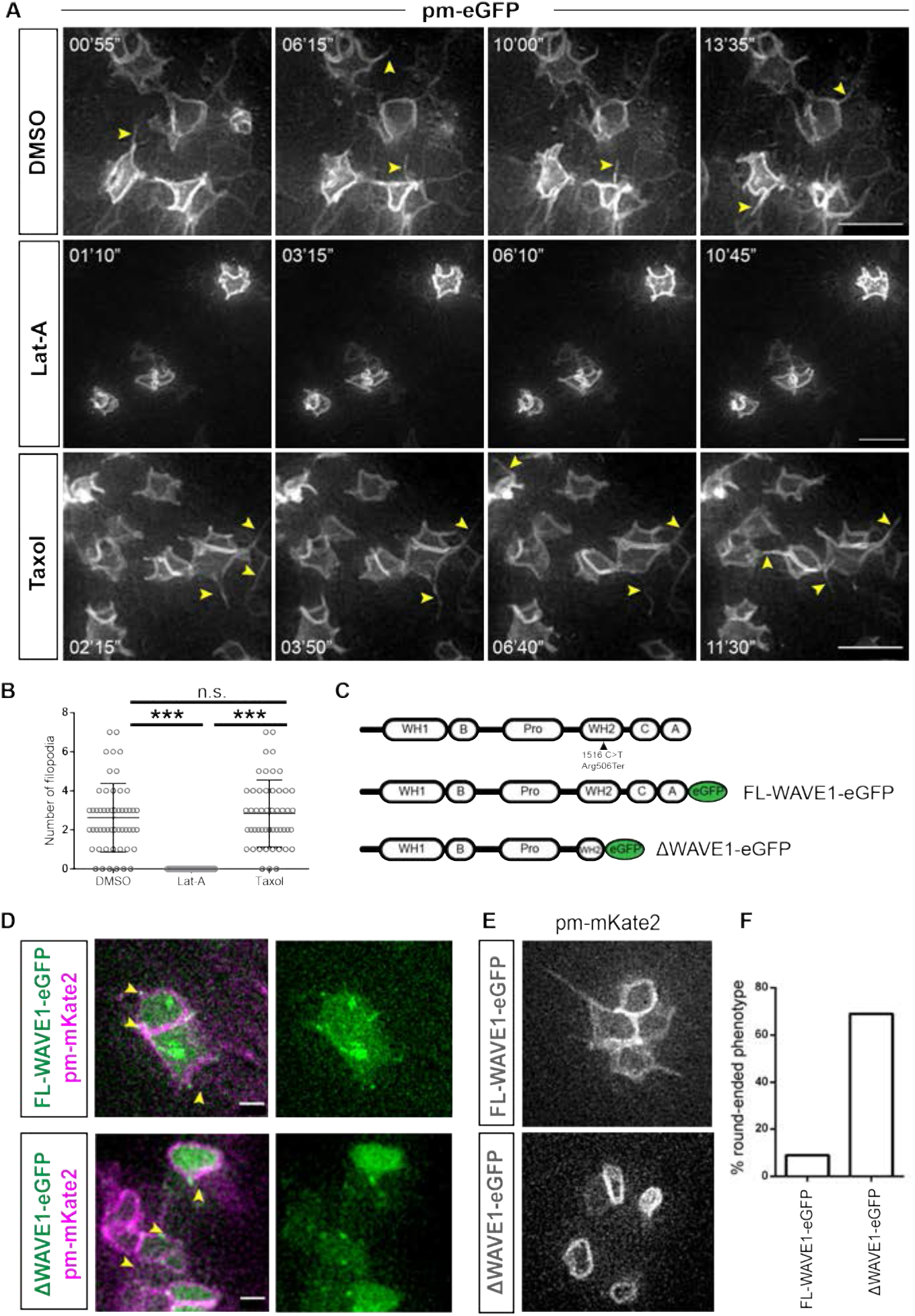
Lateral protrusion dependence on Actin/WAVE1 and ΔWAVE1 round-ended apical endfoot phenotype. (A) Live imaging timepoints monitoring lateral protrusions and filopodia in the presence of DMSO control, Latrunculin-A (Lat A) or Taxol; (B) quantification of filopodial protrusions in each condition in A, Lat A inhibits filapodial extensions compared to DMSO control and Taxol treatments, One-way ANOVA test (Tukey’s multiple comparisons test), p= 0.3852; ***p<0.0001.), error bars = mean with SD; DMSO: 3 experiments, 6 explants, 44 cells; Latrunculin-A: 3 experiments, 9 explants, 337 cells; Taxol: 2 experiments, 5 explants, 50 cells; (C) Schematic of domain composition of the human WAVE1/WASF1 protein and the nonsense mutation (Arg506Ter) identified in (Ito et al., 2018), and details of constructs used for experiments: full length (FL-WAVE1) and truncated protein (ΔWAVE1) with C-terminus eGFP tag; (D) Low-level mis-expression of FL-WAVE1-eGFP and ΔWAVE1-eGFP both localised in cell membrane at the base of lateral protrusions; (E) Round-ended apical endfoot phenotype in cells mis-expressing ΔWAVE1-eGFP, but not FL-WAVE1-eGFP, images extracted from movies 26 and 27; (F) percentage of cells with round-ended phenotype following FL-WAVE1-eGFP and ΔWAVE1-eGFP mis-expressions. FL-WAVE1-eGFP: 12/133 [9%], 4 experiments, 8 slices, 133 cells and ΔWAVE1-eGFP: 116/167 [69%], 2 experiments, 4 slices, 167 cells. Scale bars in (A) 5 μm and in (D) 2 μm.

### Lateral protrusions rely on the actin binding protein WAVE1

To monitor and manipulate actin dynamics on a cell-by-cell basis and more specifically in lateral protrusions we looked for an actin regulatory protein that might act in these structures. Branched actin filament networks are assembled through the combined activities of the Arp2/3 complex and different forms of WASP/WAVE proteins. In *Drosophila*, loss of SCAR/WAVE1 complex activity in notum epithelium results in specific loss of basolateral protrusions (Georgiou and Baum, 2010). TIRF imaging has also shown that in comparison with WAVE2 and N-WASP, WAVE1 generates shorter, slower growing actin filaments (Miki et al., 1998; Sweeney et al., 2015; Yamazaki et al., 2005), consistent with the form and dynamics of lateral protrusions we observe here. Moreover, mutations in human WASF1/WAVE1 Arp2/3/actin binding domain led to inhibition of fibroblast lamellipodial protrusion (Ito et al., 2018), further identifying WAVE1 as a potential mediator of lateral protrusions. As this region is highly conserved across species (Ito et al., 2018), we cloned and eGFP-tagged the full-length (FL) human WAVE1 and a C-terminal truncated form of the protein (informed by the nonsense mutation described by Ito et al (Figure 7C). This truncation generates a protein lacking the C and A domains and part of the WH2 domain necessary for binding actin and the Arp2/3 complex to promote actin polymerization (Boczkowska et al., 2014; Ito et al., 2018). We predicted that mis-expression of truncated (Δ) WAVE1-eGFP would have a dominant negative action, competing with endogenous WAVE complex-binding partners.

When mis-expressed at low levels, FL-WAVE1-eGFP or ΔWAVE1-eGFP localized at cell-cell junctions and at the base of lateral protrusions (Figure 7D). When ΔWAVE1-eGFP was expressed, most lateral protrusions were lost (some transient filopodia were observed, Movie 27) and the endfoot acquired a round-ended appearance: in contrast, this phenotype was only occasionally observed following high level WAVE1-eGFP mis-expression (Figures 7E, F, movies 26, 27). These data show that apical endfoot shape and lateral protrusion dynamics rely on WAVE1 and moreover, indicate that mis-expression of the ΔWAVE1 can be used to interfere with lateral protrusion formation on a cell-by-cell basis.

### Loss of lateral protrusions does not alter expression of adherens junction protein N-cadherin nor disrupt neuroepithelial tissue integrity

To address the functional significance of lateral protrusions, we first investigated whether their loss affected maintenance of adjacent adherens junctions (AJ). Adherens junction constituent protein N-Cadherin localisation appeared unaltered at the apical surface across the dorsoventral axis of the neural tube following ΔWASF1-eGFP mis-expression, suggesting that AJs remain intact, despite the loss of lateral protrusions (Figure 8A). The developing, neuron-generating, neuroepithelium lacks tight junctions which are initially located subapically, below the adherens junction (Aaku-Saraste et al., 1996). This raised the possibility that sub-apical protrusions facilitate neuroepithelial integrity. To test this 70kDa TxRd-Dextran was injected into the neural tube lumen and dextran leakage into the neuroepithelium in the presence of ΔWASF1-eGFP or control pm-eGFP was assessed; however, no differences were apparent after 24h Supplementary Figure 4). These data indicate that within this developmental window loss of lateral protrusions does not alter AJs between cells nor impact neuroepithelial tissue integrity.

**Figure 8.**
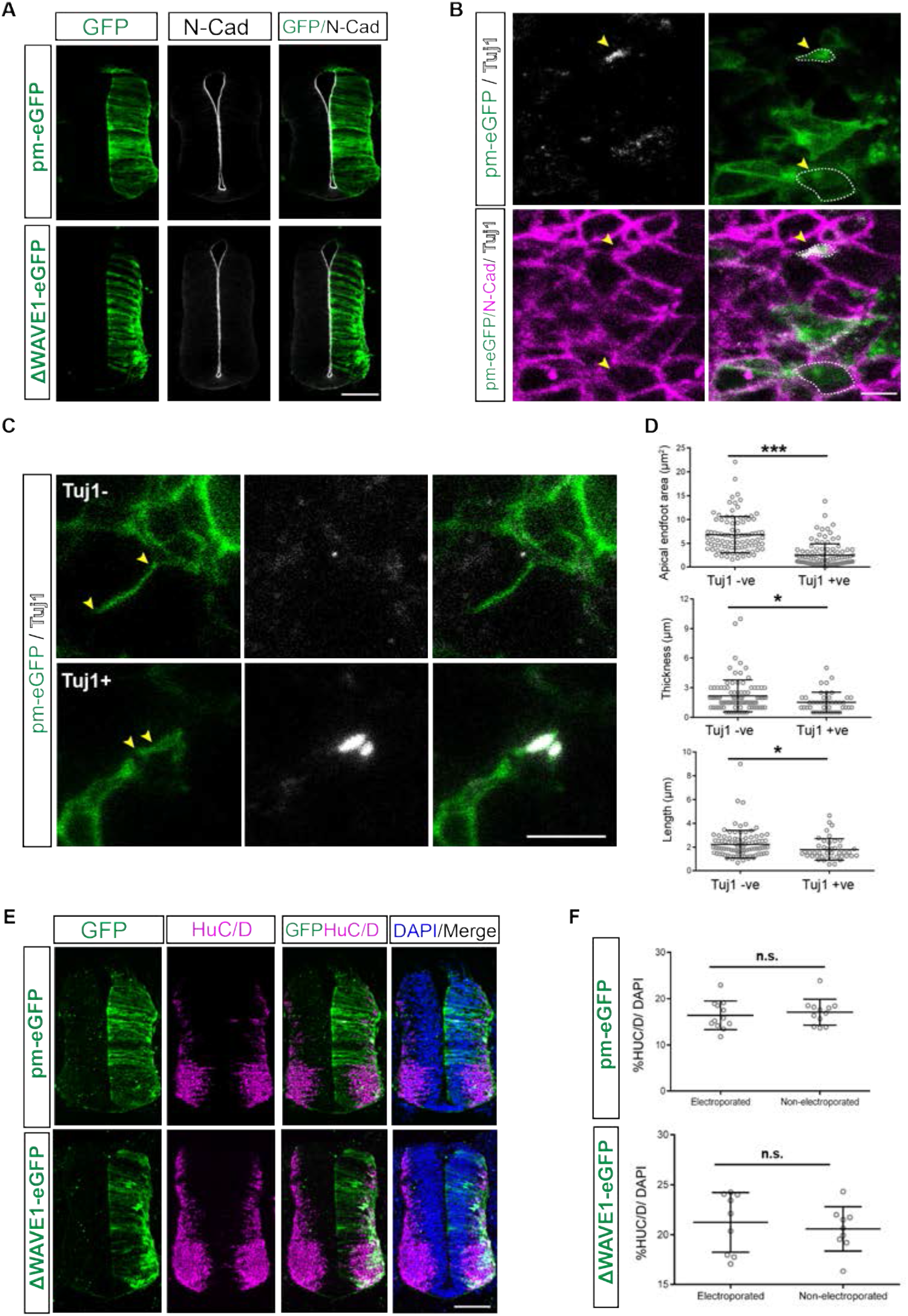
ΔWAVE1 mis-expression does not alter N-Cadherin expression and although neurogenesis involves lateral protrusion withdrawal, ΔWAVE1 does not trigger neurogenesis. (A) No alternation in AJ protein N-Cadherin following mis-expression of control pm-eGFP or ΔWAVE1-eGFP in neural tube for 48 hours (> 3 transverse sections from each of pm-eGFP = 4 embryos, ΔWASF1-eGFP = 4 embryos); (B) Tuj1+ and Tuj1-apical endfeet viewed enface at the level AJ/N-Cadherin expression (pink), white dashed lines examples of areas measured; (C) Lateral protrusion length measurement (between yellow arrowheads) in Tuj1+ and Tuj1-cells (see movies 28, 29); (D) Dot plots showing significant difference in apical endfoot area, and lateral protrusion thickness and length between Tuj1- and Tuj1+ cells (t-tests for area: 4 experiments, 10 explants, Tuj1+ cells = 103; Tuj1-cells = 93, p = 0.0001; thickness and length: 4 experiments, Tuj1+ cells = 43, Tuj1-cells = 93, thickness p = 0.0172, and length: p = 0.0335); (E) No alternation in numbers of neurons (HuC/D +ve cells) following mis-expression of control pm-eGFP or ΔWAVE1-eGFP in the neural tube for 48 hours, quantified in (F) percentage of HuC/D +ve cells on electroporated and non-electroporated sides of the neural tube (> 3 transverse sections from each of pm-eGFP = 3 embryos, ΔWASF1-eGFP = 4 embryos, t-test pm-eGFP p = 0.7496; ΔWASF1-eGFP, p = 0.4266). (C and E) Mean with standard deviation. Scale bars, A and E 100μm, C 5 μm.

### Neuronal differentiation involves withdrawal of lateral protrusions, but precocious loss of these protrusions is insufficient to trigger neurogenesis

Neuronal differentiation involves the delamination of newborn neurons, during which they are released from apical connection with neighbouring cells (Itoh et al., 2013; Kasioulis and Storey, 2018; Rousso et al., 2012; Zhang et al., 2010). This includes an abscission process which operates at the level of the AJs (Das and Storey, 2014) and may also involve withdrawal of the lateral protrusions described here. To investigate whether this latter step is involved in the regulation of neurogenesis, we first compared protrusion characteristics in TuJ1 expressing newborn neurons (which have yet to delaminate) and neural progenitors. To increase the number of newborn neurons we drove this process by mis-expressing the proneural factor neurogenin2 (Neurog2-IRES-GFP) together with pm-eGFP in the neural tube (as above). Protrusion dimensions were measured in fixed tissue enface following immunofluorescent detection of TuJ-1 and N-Cadherin for AJs. TuJ-1 expressing newborn neurons had a smaller apical endfoot area (measured at the level of the AJ) and exhibited significantly reduced lateral protrusion thickness and length (Figures 8B -D, Movies 29 – 30). These data suggest that differentiating neurons normally withdraw their lateral protrusions prior to delamination.

This finding raised the possibility that ΔWAVE1-eGFP expressing cells lacking lateral protrusions might undergo precocious neuronal differentiation. To test this, ΔWAVE1-eGFP or control pm-eGFP was mis-expressed in the neural tube and assessed after 48h. Cells expressing the neuronal marker HuC/D were then quantified, however, the proportion of HuC/D expressing cells was unaltered between electroporated and non-electroporated sides in ΔWAVE1-eGFP expressing as well as in control embryos (Figures 8E – F). Furthermore, analysis of expression of newborn neuron marker, the Notch ligand Delta1 (Dll1) (Henrique et al., 1995), in ΔWASF1-eGFP transfected embryos also revealed no changes in expression pattern at 24h (data not shown). Together, these findings suggest that while withdrawal of lateral protrusions is regulated by the neuronal differentiation programme, this step alone is insufficient to trigger this process.

## Discussion

This study provides a comprehensive live view of cellular protrusions that extend from the apical neuroepithelium and uncovers a sub-apical latticework of lateral protrusions and associated fine filopodia which mediate membrane contacts between non-neighbouring cells within this tissue. The presence of growing microtubules as well as actin in lateral protrusions, the localised expression of ENA and VASP proteins and lack of atypical Myosin-10 in these structures, together with their lamellipodia-like morphology, distinguished these structures from cytonemes. The absence of mitochondria, along with the overlap rather than fusion between lateral protrusions from distant cells further discriminate these structures from tunnelling nanotubes. Despite the presence of microtubules, lateral protrusions, and their extending filopodia, were only dependent on actin polymerization. We further show that the actin nucleating protein WAVE1 localises to lateral protrusions and that introduction of mutant WAVE1 leads to their loss. Importantly, this did not alter localisation of adherens junction protein N-cadherin nor disrupt neuroepithelial tissue integrity. While lateral protrusions were specifically withdrawn prior to delamination of newborn neurons, experimentally induced precocious loss of these structures was insufficient to trigger neurogenesis. These findings suggest that lateral protrusion withdrawal is regulated by the neuronal differentiation programme and that re-configuration of the actin/WAVE1 network is an early step that anticipates neuronal delamination. Overall, this study identifies a dynamic sub-apical latticework of lateral protrusions which has the potential to mediate cell-cell communication across the developing neuroepithelium.

Ultra-structure analyses and live imaging of the lumen facing apical membrane of neuroepithelial cells revealed three distinct apically protrusive structures, in addition to the primary cilium; microvilli (also known as micro-spikes), lamellopodia-like structures and long filopodia, all of which could be further visualized with F-tractin and so are actin-based. The presence of microspikes on neuroepithelial cells has been reported previously using electron microscopy to analyze the closing neural tube in chicken and rodent embryos (Shepard et al., 1997, 1993; Smith et al., 1982; Waterman, 1976). Later work revealed localization of the surface glycoprotein prominin-1 (CD133) on neuroepithelial cell microvilli (Marzesco et al., 2005), suggesting that these serve to increase the apical surface area for internalization of potential growth factors present in the lumen. Most recently, in Caco-2 intestinal epithelial cells, actin assembly has been shown to drive microspike protrusions toward apical adherens junctions and increase the surface membrane available for Cadherin mediated cell-cell-adhesion (Li et al., 2021, 2020) which might be conserved in the neuroepithelium. We further document, to our knowledge, novel apical lamellipodial protrusions, the dimensions of which suggest that they can reach over neighbouring apical endfeet and so may influence their neighbours and/or operate to sense the immediate environment. Finally, we observed long apical filopodia that contact non-neighbouring cells. Such protrusions have been noted previously in fixed tissue (Shepard et al., 1997, 1993; Waterman, 1976). We reveal here that these apical filopodia exhibit retrograde membrane movement indicative of active endocytosis of components from the environment. In support of this possibility, signaling pathways such as sonic hedgehog and Wnt show the binding and trafficking of the ligands along such cytoneme-like membrane protrusions in the chick limb mesenchyme (Sanders et al., 2013) and early zebrafish neural plate (Stanganello et al., 2015).

The presence of lateral protrusive structures has been reported previously in a range of non-neural epithelia (Demontis and Dahmann, 2007; Farquhar and Palade, 1965; Tang and Brieher, 2012). Recent work in developing mouse cerebral cortex identified very similar lateral lamellipodia-like structures by EM and reported their actin dependence (Shinoda et al., 2018). We confirm the sub-apical emergence of lateral protrusions in the chicken and human spinal cord using EM. In the chick, live imaging further revealed that these lamellipodia-like protrusions create contacts between non-neighbouring cells and provide a platform for filopodial extensions, forming dynamic membrane contacts with a radius of at least two apical endfeet (> ~5 microns) from each neuroepithelial cell: effectively forming a latticework of cell contacts within the neuroepithelium. Our findings suggest that these contacts do not involve cell fusion (they extend along each other and appear not to traffic mitochondria) and so are distinct from tunnelling nanotubes (Korenkova et al., 2020; Rustom et al., 2004). However, future studies could assess further potential trafficked organelles, including lysosomes and endosomes.

The lamellipodial-like structure of lateral protrusions shares some similarity with basal protrusions described in *Drosophila* notum epithelium (Georgiou and Baum, 2010). Importantly, these basal protrusions are uniquely SCAR/WAVE dependent and consistent with this we found the related branched actin nucleating protein WAVE1 (Carlier et al., 2015; Pollard and Borisy, 2003) was localised to and required for lateral protrusions. Moreover, WAVE regulatory complexes work cooperatively with Ena/VASP to enhance Arp2/3 mediated actin assembly in lamellipodia (Chen et al., 2014) and see (Damiano-Guercio et al., 2020), and ENA and VASP localisation to lateral protrusion tips here further identifies these as lamellipodia-like structures. However, it is important to note that WAVE1 also localises to mitochondria in fibroblasts (Ito et al., 2018), and we observe this in neuroepithelial cells too, so its regulation of lateral protrusions could also reflect an indirect WAVE1 function in this organelle. Lateral protrusions were additionally distinguished by the presence of polymerising microtubules from base to tip, while most cellular protrusions are exclusively composed of actin, or have microtubules confined to the base (González-Méndez et al., 2019). Indeed, together these findings identify a surprising similarity between sub-apical lateral protrusions and their extending filopodia and the cytoskeletal organisation of later forming axonal growth cones (Robert Manak, 2018).

The discovery that lateral protrusions are withdrawn by newborn neurons that are still attached at the apical surface suggests that this step is an early event directed by the neuronal differentiation programme. Moreover, precocious reduction of lateral protrusions induced by mis-expression of a mutant form of WAVE1 was insufficient to trigger neurogenesis. Importantly, neuronal delamination and differentiation is elicited by loss of AJ constituent proteins N or E Cadherin (Hatakeyama et al., 2014; Itoh et al., 2013; Rousso et al., 2012). It is therefore not surprising that reduction of WAVE1 and lateral protrusive activity also did not impact N-Cadherin expression or localization. This is further consistent with findings in the *Drosophila* notum epithelium, where loss of SCAR/WAVE1 complex activity similarly does not impact apico-basal polarity (Georgiou and Baum, 2010). These findings suggest that regulation of WAVE1 activity in lateral protrusions is an early action of the neuronal delamination mechanism and supports the notion that this anticipates the extensive actin network re-configuration apparent during apical abscission (Das and Storey, 2014; Kasioulis et al., 2017). The importance of actin configuration in this process is further underlined by the requirement for the actin cross-linking protein Filamin A, mutation of which underlies defective neuronal delamination in the human condition periventricular heterotopia (Parrini et al., 2016; Sheen et al., 2001; Zhang et al., 2013).

Overall, our findings indicate that withdrawal of lateral protrusions is regulated by, but does not drive, the neuronal differentiation programme and that these structures may be part of the mechanism that apically anchors cells within the neuroepithelium. However, this does not exclude roles in cell communication for this latticework of cell membrane contacts (Figure 9). One possibility is that lateral protrusions, which radially contact cells several endfeet away, facilitate cell-cell coupling within cell micro-clusters, which have recently been shown to exhibit synchronous Hes gene (Notch signalling) oscillations, and so may serve to regulate neuron production (Biga et al., 2021). Indeed, a role in local coordination of cell signalling is consistent with lateral protrusion withdrawal in newborn neurons, which might then experience reduced signalling from neighbouring cells, while subsequently delivering Notch ligand via adherens junctions (Hatakeyama et al., 2014), and see (Falo-Sanjuan and Bray, 2021). This an interesting idea because it could provide a cell biological basis for the compartmentalization of cell selection within the developing neuroepithelium. The lateral protrusion latticework also provides an opportunity to relay signals across the developing neuroepithelium. Such relays may be required to convey signalling beyond the reach of ligand diffusion in extracellular space/neural tube lumen and/or delivery by cytonemes/long filopodia. The potential for lateral protrusions to facilitate comparison and integration of signalling between cells in this further context suggests a role in establishing and/or maintaining positional information across the neuroepithelium. To elucidate these functional roles, future studies should assess local patterns of contact dynamics and localise ligands and receptors in lateral protrusions.

**Figure 9.**
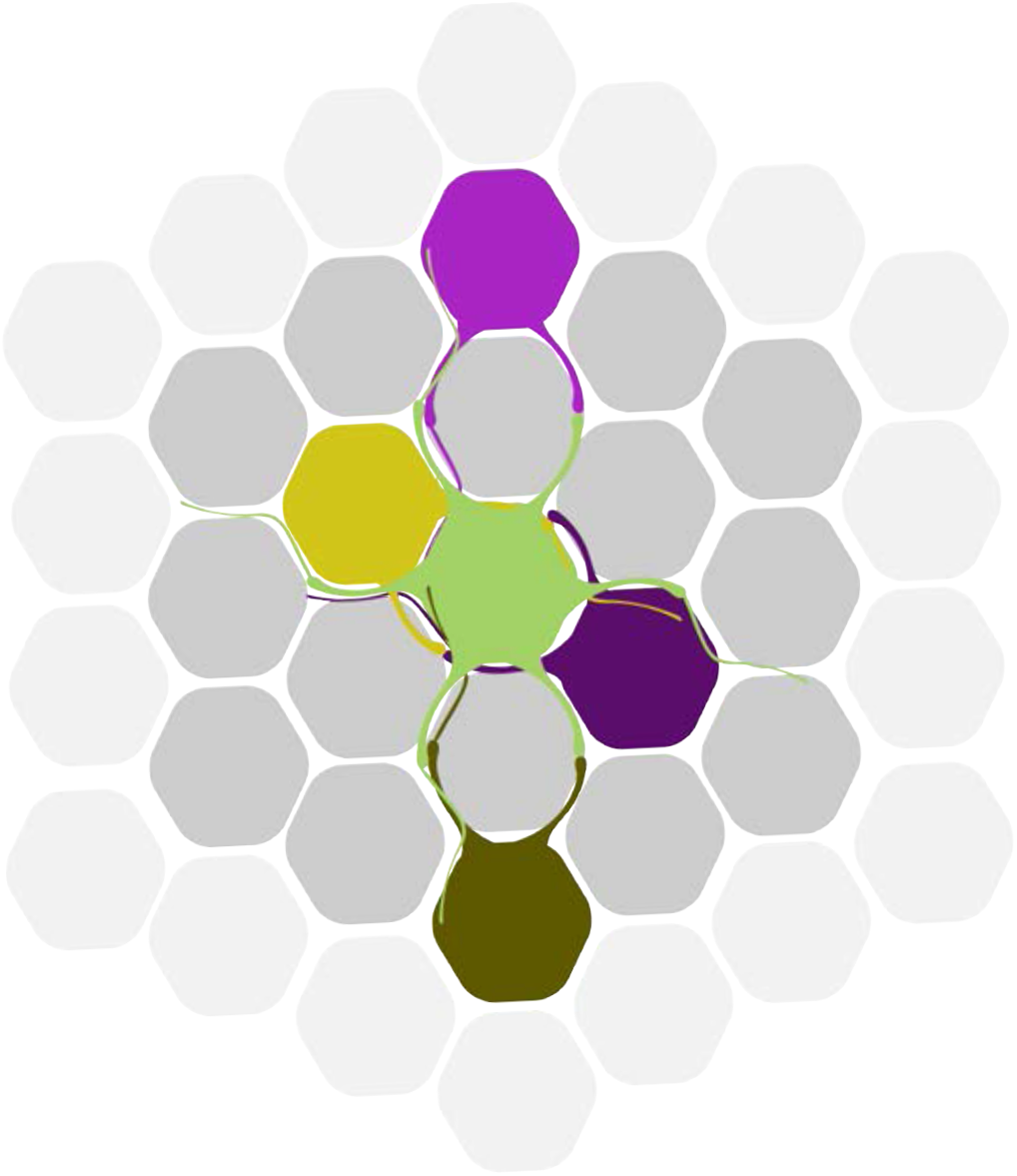
The lateral protrusion latticework. Schematic depicting exemplar interactions of lateral protrusions extending from one apical end foot (light green). Lateral protrusions with filipodia can make contacts with cells and their protrusions 1-2 endfeet away (light purple and dark green), while also experiencing mutual contacts with their immediate neighbours (yellow, dark purple). Lateral protrusions from non-neighbouring cells can meet and retract or extend along each other. As all apical endfeet extend lateral protrusions this creates a latticework of short-range cellular protrusions with the potential to mediate cell-cell communication across the neuroepithelium.

## Supporting information

Movies 1-29 Kasioulis et al

## Acknowledgements

We are grateful to our colleagues Kees Weijer and Jens Januschke for comments on this manuscript, to Raman Das for preparation of chicken embryonic spinal cord for electron microscopy and imaging by John James presented in Figures 1A-C and to Professor Tim Sanders, (The Neuroscience Institute University of Chicago), for sharing plasmid constructs. Human embryonic tissue in this study was provided by a Joint MRC/Wellcome Trust grant (# MR/R006237/1) to the Human Developmental Biology Resource (www.hdbr.org). IK, AD and PH were supported by a Wellcome Investigator award to KGS (WT102817AIA). The confocal microscope used for imaging was purchased with support from a Wellcome Trust Multi-User Equipment grant (WT101468).

The authors declare that they have no conflict of interest.

## Materials and Methods

### Tissue sources

Fertilised Bovans Brown chicken eggs were supplied by Henry Stewart & Co. Ltd and incubated at 38 °C to the desired Hamburger and Hamilton (HH) stage. Work with chicken embryos at the stages investigated here do not require ethical approval. Fixed human embryonic spinal cord tissue at Carnegie stage (CS)12 used for EM studies was provided by the Human Developmental Biology Resource (HDBR) project # 200407.

### Cloning

The ImProm-II reverse transcription system (Promega, A3800) was used to generate cDNA from hESC (H9) RNA. Full length WAVE1 was PCR amplified adding in a linker sequence. The WAVE1 PCR product was then used to amplify the mutant WAVE1 (ΔWAVE1), (a truncated version of the gene lacking sequence distal to nonsense mutation Arg506Ter, Ito et al 2018, see Figure 6C) to which the linker sequence was added in a second round of PCR. Both were tagged with monomeric eGFP, separated by the linker sequence, in a further PRC reaction. For primers for full length WAVE1 (FL) and ΔWAVE1 see Table 1. PCR reactions were carried out using the Phusion polymerase (ThermoFisher, F537). Following each reaction, the products were gel extracted using the Zymoclean Gel DNA recovery kit (D4001). The pENTR/D-TOPO cloning kit (ThermoFisher, K240020) was used to introduce the PCR products into the entry vector for Gateway cloning. PCR products in the entry vector were then recombined in the destination vector using the LR clonase II enzyme mix (ThermoFisher, 11791100). We utilised the PBX destination vector (kind gift from Timothy Sanders (Sanders et al., 2013) which contains the CAG promoter, for efficient expression in chicken tissue.

**Table 1.**
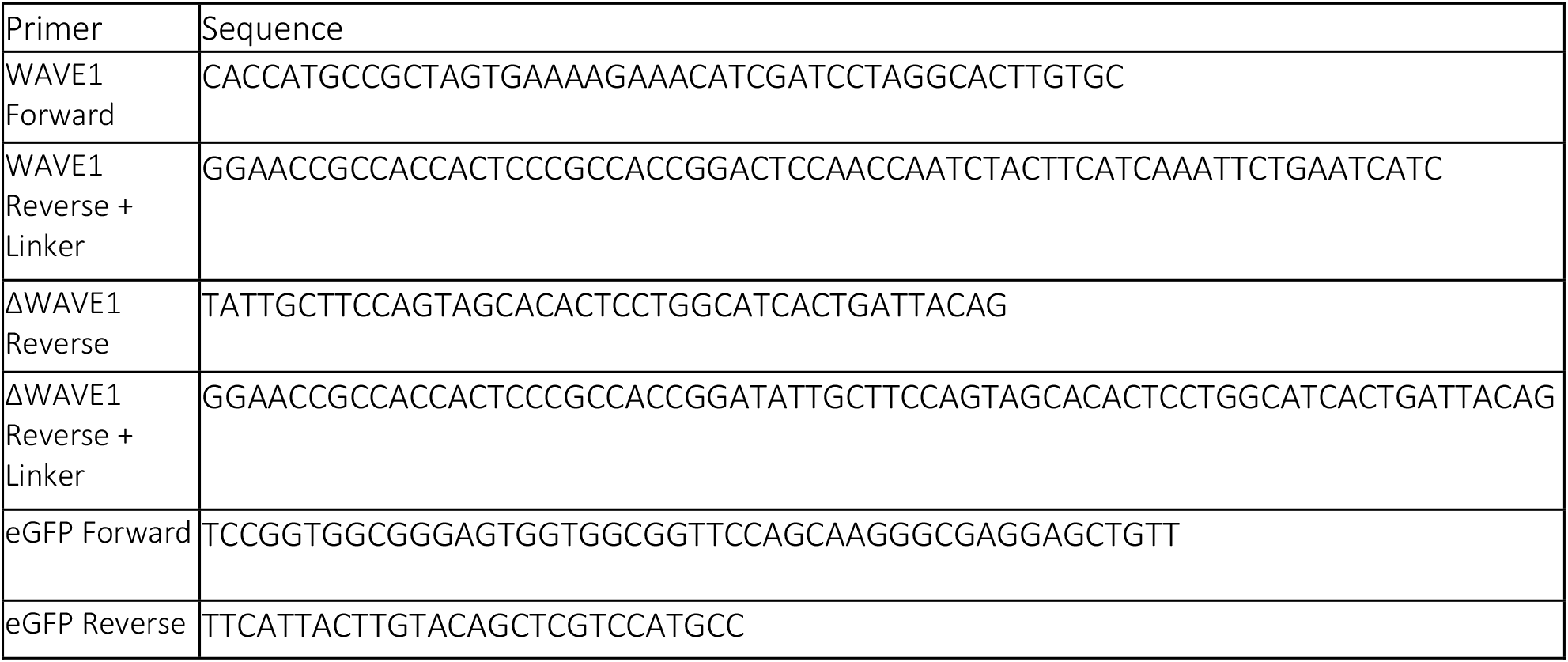
Primer sequences.

### In ovo electroporation

Fertilised chicken eggs were incubated at 38 °C to stage HH 10-12 and the neural tube was electroporated with a single or combination of constructs at concentrations given in Table 2. We targeted the neural tube region posterior to the forelimb level and along the dorso-ventral axis, transfected cells were typically located throughout the dorsal region and extended to include the Olig2-positive ventral domain. To obtain mosaic expression pattern, neural tubes were electroporated with three 50ms pulses, separated between them with a 950ms gap, at 17 Volts using the ECM 830 square wave electroporation system (BTX).

**Table 2.** Plasmid constructs, source and dilution.

| Plasmid | Source | Dilution |
| --- | --- | --- |
| pm-eGFP | Tim Sanders, University of Chicago | 50 – 100 ng/μl |
| pm-mKate2 | Tim Sanders, University of Chicago | 50 – 100 ng/μl |
| F-tractin-mKate2 | Tim Sanders, University of Chicago | 50 ng/μl |
| EB3-sfGFP | Tim Sanders, University of Chicago | 25 ng/μl |
| WAVE1-eGFP | See cloning protocol | 1 μg/μl |
| ΔWAVE1-eGFP | See cloning protocol | 1 μg/μl |
| MyoX-eGFP | Tim Sanders, University of Chicago | 1 μg/μl |
| mNeonGreen-ENA | Allele Biotechnology | 1 μg/μl |
| mNeonGreen-Vasp | Allele Biotechnology | 1 μg/μl |
| mNeonGreen-TOMM20 | Allele Biotechnology | 1 μg/μl |

### Embryonic tissue explant culture

Electroporated embryos were allowed to develop overnight to stage HH17-18 and the neural tube in the interlimb region was then bisected along the dorsovental axis, to gain access to the neuroepithelial apical surface. The non-electroporated side was discarded and hemi neural tube explants (~ 2-3 somite lengths), including associated somites, were embedded for enface imaging, with the neuroepithelial apical side orientated to face the base of the culture dish. In most experiments only one explant per embryo was used. Explants were embedded in collagen (supplemented with 0.1% acetic acid, 5x L-15 medium [ThermoFisher, 41300] and 7.5% sodium bicarbonate [ThermoFisher, 25080094]) in poly-D-lysine coated glass-bottomed petri-dishes (World Precision Instruments, FD35-PDL- 100), covered with neurobasal media, supplemented with B-27 (ThermoFisher, 17504044), glutamax (ThermoFisher, 35050038) and gentamicin (ThermoFisher, 15750037). The plates were then placed in a 37°C-maintained incubator supplied with 5% CO_2_, giving time for the explants to recover for two hours before commencing live imaging (Das et al., 2012).

### Time lapse imaging

Time-lapse imaging of neural tube explants was performed using a Deltavision Core microscope system in a WeatherStation environmental chamber maintained at 37°C. (GE Healthcare). Imaging was optimised for minimal light transmission, exposure times (50–100 milliseconds) and total acquisition times of 10-20 minutes, to avoid phototoxicity (Das et al., 2012; Das and Storey, 2014). Image acquisition was performed using an Olympus 60 × 1.42 NA oil immersion objective or an Olympus 40 × 1.25 NA silicone oil immersion objective, a solid-state LED light source and a CoolSnap HQ2 cooled CCD camera (Photometrics). Unless otherwise stated, 10 optical sections spaced 0.5 μm apart were acquired for each explant (1024 × 1024 pixels, 1 × 1 binning). For the live imaging sessions totalling for 1 hour, image acquisition between timepoints was set at 1-minute intervals. Images were deconvolved using the SoftWorx image processing software.

### Preparation of tissue for imaging cytoskeletal manipulations

To investigate the effect of actin polymerisation inhibition or microtubule stabilisation, neural tube explants embedded for enface imaging were incubated in pre-warmed neurobasal medium containing latrunculin-A (1 μM, Abcam, ab144290) or Taxol (10 μM, Sigma, T7191) for 1 hour before imaging.

### Confocal imaging and 3D modelling

Imaging of sections or embedded tissue (Figures 8A, B & D) was carried out with the Leica SP8 confocal microscope (lenses HC PL APO CS2 20x/0.75 DRY or HC PL APO CS2 63x/1.40 OIL). 3D modelling for Movie 7 was carried out using the IMARIS software (Oxford instruments).

### Immunofluorescence and fixed tissue imaging

The interlimb region of HH17–18 chick embryos was fixed in 4% paraformaldehyde for 2 hours at 4°C, washed with PBS and equilibrated overnight in 30% sucrose at 4°C. The tissue was then embedded in 1.5% LB agar (Sigma, L7025) and 5% sucrose dissolved in PBS. Mounted tissue was dehydrated for 48 hours in 30% sucrose and snap frozen on dry ice. 20-25 μm thick transverse sections were then collected using a Leica cryostat (maintained at −25°C). For immunofluorescence experiments, sections were re-hydrated in 0.1% Triton-X-100/PBS, incubated in blocking buffer for 1-2 hours at room temperature (0.1% Triton-X-100% and 1% heat inactivated donkey and goat serum, in PBS) and incubated overnight with primary antibodies (diluted in blocking buffer) as in Table 3. Sections were washed with 0.1% Triton-X-100/PBS and incubated overnight with secondary antibodies for 2 hours at room temperature. Following washes with 0.1% Triton-X-100/PBS, the sections were mounted in Prolong Gold antifade mountant (ThermoFisher, P36930). For enface imaging of fixed tissue, explants were mounted in 0.5% low gelling temperature agarose (Sigma, A9045). For enface confocal imaging following a live imaging session, the neurobasal medium was immediately replaced with 4% paraformaldehyde for ~1 hour, washed and maintained in PBS. DAPI was added in the PBS washes to stain the nuclei.

**Table 3.** Antibodies, stains, sources and dilutions.

| Antibody | Source | Catalogue number | Dilution |
| --- | --- | --- | --- |
| **Primary antibodies** |  |  |  |
| Rat N-Cadherin | ThermoFisher | 132100 | 1/300 |
| Tuj1 | Biolegend | 801213 | 1/500 |
| Mouse HuC/HuD | ThermoFisher | A21271 | 1/200 |
| Goat GFP | Abcam | 6673 | 1/500 |
| Sheep Dll1 | R&D | AF3970 | 1/75 |
| Donkey anti-rat 568 | Abcam | ab175475 | 1/300 |
| **Secondary antibodies** |  |  |  |
| Donkey anti- mouse 568 | ThermoFisher | A-10037 | 1/300 |
| Donkey anti- mouse 647 | ThermoFisher | A-31571 | 1/300 |
| Donkey anti-goat 488 | ThermoFisher | A-11055 | 1/300 |
| Donkey anti-sheep 488 | ThermoFisher | A-11015 | 1/300 |
| Phalloidin CF640R | Biotum | 00050 | 1/300 |

### EM sample preparation

Chicken embryonic spinal cord at HH12 was fixed in 1% paraformaldehyde and 1.5% glutaraldehyde in 0.05 M cacodylate buffer (pH 7.0) for two hours at room temperature and was post-fixed with 1% osmium tetroxide (OsO4). After washing in cacodylate buffer the tissue was stained with 1% tannic acid in 0.1M cacodylate buffer for 1hr. The tissue was then dehydrated through an alcohol series finishing in propylene oxide. The tissue was then infiltrated with Durcupan epoxy resin (Sigma), sectioned with a Leica UCT ultramicrotome and imaged on a FEI Tecnai TEM using the SIS Megaview II CCD camera (Figures 1A-C). Human embryonic tissue fixed in 4% paraformaldehyde and 2.5% glutaraldehyde in 0.1M sodium cacodylate buffer (pH 7.2) for 60 minutes was provided by the HDBR. The tissue was washed twice in cacodylate buffer, and small pieces of spinal cord were then post-fixed in 1% OsO4 with 1.5% Na ferricyanide in cacodylate buffer for 60 min. After another cacodylate buffer wash they were contrasted with 1% tannic acid and 1% uranyl acetate. The cell pellets were then serially dehydrated into 100% ethanol, changed to propylene oxide left overnight in 50% propylene oxide 50% resin and finally embedded in 100% Durcupan resin (Sigma). The resin was polymerised at 60°C for 48hrs and sectioned on a Leica UCT ultramicrotome. Sections were contrasted with 3% aq uranyl acetate and Reynolds lead citrate before imaging on a JEOL 1200EX TEM using a SIS III camera. Chick embryonic tissue imaged in Supplementary figure3 was prepared using the same protocol as for human tissue.

### Tissue integrity assay using fluorescent dextran

Chick embryos were electroporated with pm-eGFP or ΔWAVE1-eGFP at HH10-12 as above and at stage HH17-18, the 70kDa TxRd-tagged Dextran, lysine fixable (ThermoFisher, D1864) (reconstituted following manufacturer’s instructions) was injected into the neural tube lumen. Embryos were incubated for an hour and then dissected and fixed immediately with cold 4% PFA, followed by gentle washing in PBS to remove the excess dextran, embedding and cryo-sectioning prior to confocal imaging (as above).

### Software for quantification and analysis

The Softworx software (Applied Precision, Inc), was used for the quantifications in Figure 2B, 2D, 3A, 3B, 3C, 3D, 3F, 4A, and supplementary figure 1B. Fiji software (Schindelin et al., 2012) was used for the quantifications in Figures 3F, 6B, 6C, 6D, 7B, 7F, 8D, 8F, supplementary figures 1A, 2A and 2C. Results of statistical analyses undertaken using Prism 8 are presented in figure legends, full data and analysis is provided in metadata files.

### Metadata files

Figure 2B_Microvilli_and_Lamellipodia.xls

Figure 2D_Surface protrusions.xls

Figure 3A-B and Supplementary 1B.xls

Figure 3C-D and Supplementary 1C

Figure 6B – D Lateral filopodia.xls

Figure 6C and 7B-Lateral filopodia quantifications.xlsx

Figure 7F_Quantification of round-ended apical endfoot phenotype.xls

Figure 8D-Lateral protrusions in differentiating Tuj1 positive cells.xlsx

Figure 8F-HuCD calculations.xlsx

Supplementary Figure 1A-Apical endfoot diameter

Supplementary Figure 2A-EB3-sfGFP_pm-mKate2.xlsx

Supplementary Figure 2B-EB3-sfGFP_F-tractin-mKate2-log-files.xlsx

Supplementary Figure 2E.xls

## Movie legends

**Movies 1, 2**: Microvilli (movie 1) and lamellipodial (movie 2) formation at the apical cell membrane. These movies are related to Figure 2A. Mis-expressed pm-eGFP labels the plasma membrane and F-tractin-mKate2 the actin cytoskeleton. Maximum intensity projections of deconvolved image sequences were used. Images acquired every 8 seconds.

**Movie 3**: Microvilli formation at the apical cell membrane. This movie is related to Figure 2A. Apico-basal Z-stack series across the apical plasma membrane, labelled with pm-eGFP, sub-apical N-Cadherin based adherens junctions (white) help define apico-basal position.

**Movie 4**: Long thin protrusions extending into the lumen exhibiting retrograde movement of membrane particles. This movie is related to Figure 1C. Mis-expressed pm-eGFP labels the plasma membrane and F-tractin-mKate2 the actin cytoskeleton. Maximum intensity projections of deconvolved image sequences were used. Images acquired every 7 seconds.

**Movie 5:** Sub-apical lateral protrusions arise beneath the N-Cadherin based adherens junctions. Apico-basal Z-stack series across the apical plasma membrane, labelled with pm-eGFP, sub-apical N-Cadherin (pink) based adherens junctions and nuclei (DAPI, blue).

**Movie 6**: Sub-apical lateral protrusion dynamics. This movie is related to Figure 2F. Mis-expressed pm-eGFP labels the plasma membrane. Maximum intensity projections of deconvolved image sequences were used. Images acquired every 5 seconds.

**Movie 7**: A variety of lateral protrusion dimensions revealed by confocal imaging and 3D models. This movie is related to Supplementary Figure 1A. On the left, apico-basal Z-stack series across the apical plasma membrane and on the right a 3D model projection from the former imaging. Mis-expressed pm-eGFP labels the plasma membrane.

**Movie 8, 9**: Contact of lateral protrusions from non-neighbouring apical endfeet. These movies are related to Figure 3E. Mis-expressed pm-eGFP labels the plasma membrane. Maximum intensity projections of deconvolved image sequences were used. Images acquired every 5 seconds.

**Movie 10:** Sub-apical protrusions extended around the plasma membrane of neighbouring endfeet. This movie is related to Figure 4C. Mis-expressed pm-eGFP and pm-mKate2 label the plasma membrane. Maximum intensity projections of deconvolved image sequences were used. Images acquired every 15 seconds.

**Movie 11, 12**: Lateral protrusions from non-neighbouring apical endfeet meet tip to tip and extend along each other. These movies are related to Figures 4D,E. Mis-expressed pm-eGFP and pm-mKate2 label the plasma membrane. Maximum intensity projections of deconvolved image sequences were used. Images acquired every 10 or 15 seconds.

**Movie 13**: Lateral protrusions contain actin. This movie is related to Figure 5A. Mis-expressed pm-eGFP labels the plasma membrane and F-tractin-mKate2 the actin cytoskeleton. Maximum intensity projections of deconvolved image sequences were used. Images acquired every 8 seconds.

**Movie 14:** Microtubule growth in lateral protrusions is unidirectional. This movie is related to Figure 5B. Mis-expressed pm-mKate2 labels the plasma membrane and EB3-sfGFP the polymerizing microtubules. Maximum intensity projections of deconvolved image sequences were used. Images acquired every 6 seconds.

**Movie 15, 16**: mNeonGreen-ENAH or mNeonGreen-VASP localize at adherens junctions. These movies are related to Supplementary Figure 2D. Mis-expressed F-tractin-mKate2 labels the actin cytoskeleton. Maximum intensity projections of deconvolved image sequences were used. Images acquired every 10 seconds.

**Movie 17, 18**: mNeonGreen-ENAH or mNeonGreen-VASP localize at lateral protrusions. These movies are related to Figure 5C. Mis-expressed F-tractin-mKate2 labels the actin cytoskeleton. Maximum intensity projections of deconvolved image sequences were used. Images acquired every 10 seconds.

**Movie 19:** Myosin X (MyoX-eGFP) does not localise to lateral protrusion tips. This movie is related to Supplementary Figure 2F. Mis-expressed pm-mKate2 labels the plasma membrane. Maximum intensity projections of deconvolved image sequences were used. Images acquired every 10 seconds.

**Movies 20, 21**: Thin filopodia extend from lateral protrusions. These movies are related to Figure 6A. Mis-expressed pm-eGFP labels the plasma membrane. Maximum intensity projections of deconvolved image sequences were used. Images acquired every 1 minute.

**Movie 22:** No mitochondrial transfer between cells through the lateral protrusions. This movie is related to Supplementary Figures 3A, B. Mis-expressed mNeonGreen-TOMM20 labels the mitochondria and pm-mKate2 the plasma membrane. Maximum intensity projections of deconvolved image sequences were used. Images acquired every 10 seconds.

**Movies 23-25**: Formation of lateral protrusions depend on actin but not microtubules. These movies are related to Figure 7A (movie 2, DMSO control, movie 24 LatA treated; Movie 25 Taxol treated). Mis-expressed pm-eGFP labels the plasma membrane. Maximum intensity projections of deconvolved image sequences were used. Images acquired every 5 seconds.

**Movies 26, 27**: Lateral protrusions following mis-expression of full length WAVE1-eGFP (movie 26) or mutant WAVE1-eGFP (movie 27), the latter exhibiting a round ended apical endfoot phenotype. These movies are related to Figure 7E. Mis-expressed pm-mKate2 labels the plasma membrane. Maximum intensity projections of deconvolved image sequences were used. Images acquired every 1 minute.

**Movie 28:** Tuj1 negative progenitors have long lateral protrusions. This movie is related to Figure 8B. Apico-basal Z-stack series across the apical plasma membrane, labelled with pm-eGFP and Tuj1 (left) or pm-eGFP, Tuj1 and N-Cadherin (right).

**Movie 29:** Tuj1 positive neurons have short lateral protrusions. This movie is related to Figure 8B. Apico-basal Z-stack series across the apical plasma membrane, labelled with pm-eGFP and Tuj1 (left) or pm-eGFP, Tuj1 and N-Cadherin (right).

---

**Supplementary Figure 1.**
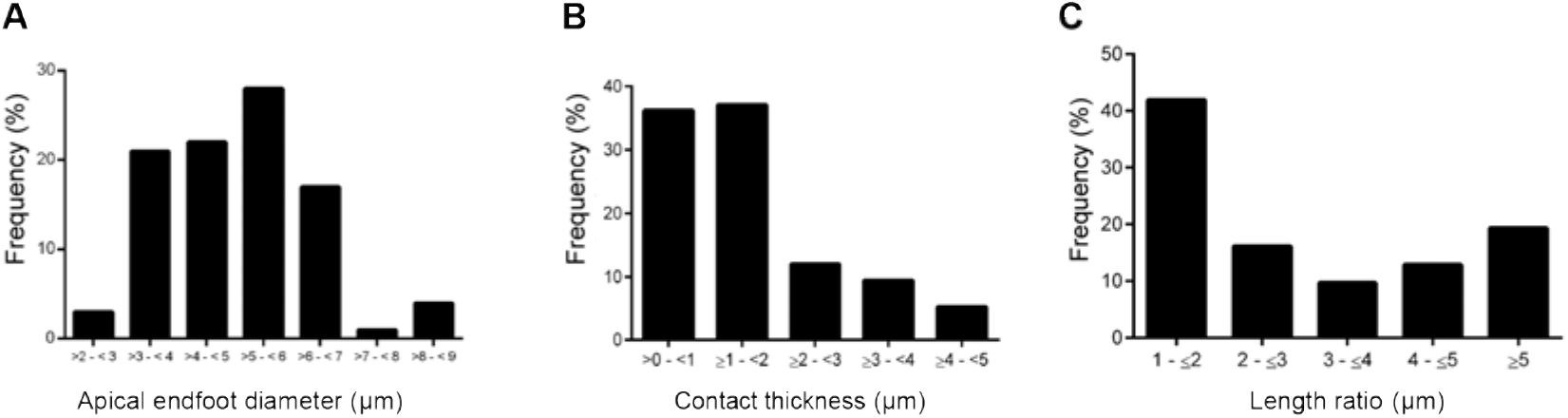
Apical endfoot and lateral protrusion contact dimensions. (A) Range of apical endfoot diameters, most are between 3 and 6 μm (5 experiments, 9 explants, 98 cells); (B) Contact protrusion thickness between non-neighbouring cells. In more than 70% of protrusion pairs, the contacting thickness did not exceed 2 μm (4 experiments, 7 explants, 116 protrusion pairs); (C) Length ratio between contacting protrusions and their frequency (4 experiments, 8 explants, 31 protrusion pairs).

**Supplementary Figure 2.**
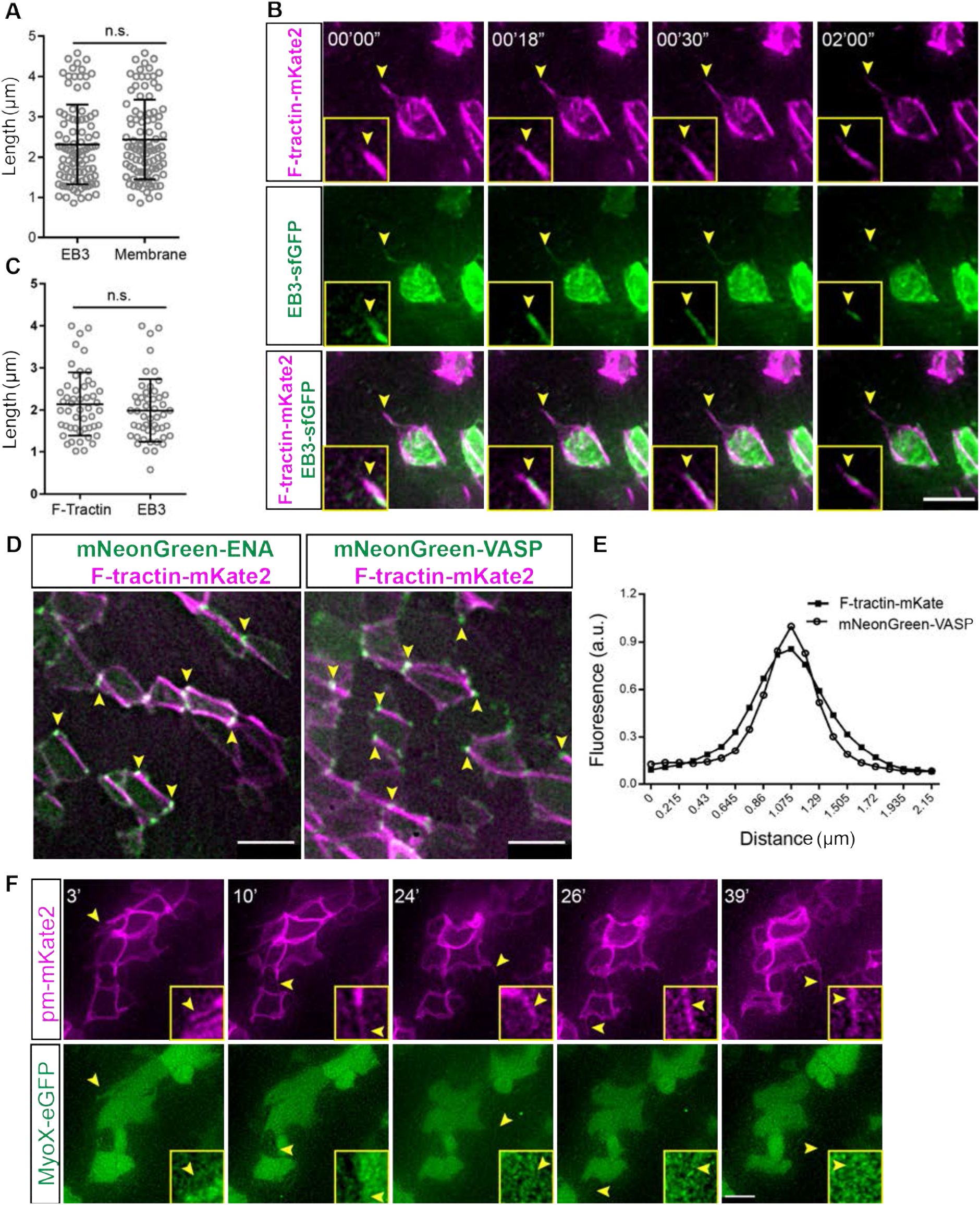
Lateral protrusions cytoskeletal architecture. (A) Dot plot for maximal length travelled by EB-sfGFP comets in protrusions and the length of lateral protrusions using the membrane marker pm-mKate2 (2 experiments, 5 explants, 93 protrusions; t-test, n.s.= not significant, p = 0.3990); (B) Live imaging timepoints following mis-expression of EB-sfGFP and pm-mKate2, arrowheads and image insets show membrane protrusion tip and EB-sfGFP localization; (C) Dot plot for maximal length travelled by EB-sfGFP comets and the length of F-tractin-mKate2 in the same protrusions (2 experiments, 6 explants, 50 protrusions; t test, n.s. = not significant, p = 0.3104); (D) Live imaging timepoints following mis-expression of mNeonGreen-ENA or mNeonGreen-VASP with F-tractin-mKate2. At the level of adherens junctions, both mNeonGreen-ENA or mNeonGreen localized at cell junctions (yellow arrowheads); (E) Line graph showing fluorescence intensity of mNeonGreen-VASP and F-tractin-mKate2 at the cell junctions (2 experiments, 5 explants, 50 measurements); (F) Live imaging timepoints for MyoX-eGFP and pm-mKate2. MyoX-eGFP was detected in cytoplasm but did not localize to lateral protrusions (arrowheads). (A, C) Mean with standard deviation. Scale bars (B, D, F) 5 μm.

**Supplementary Figure 3.**
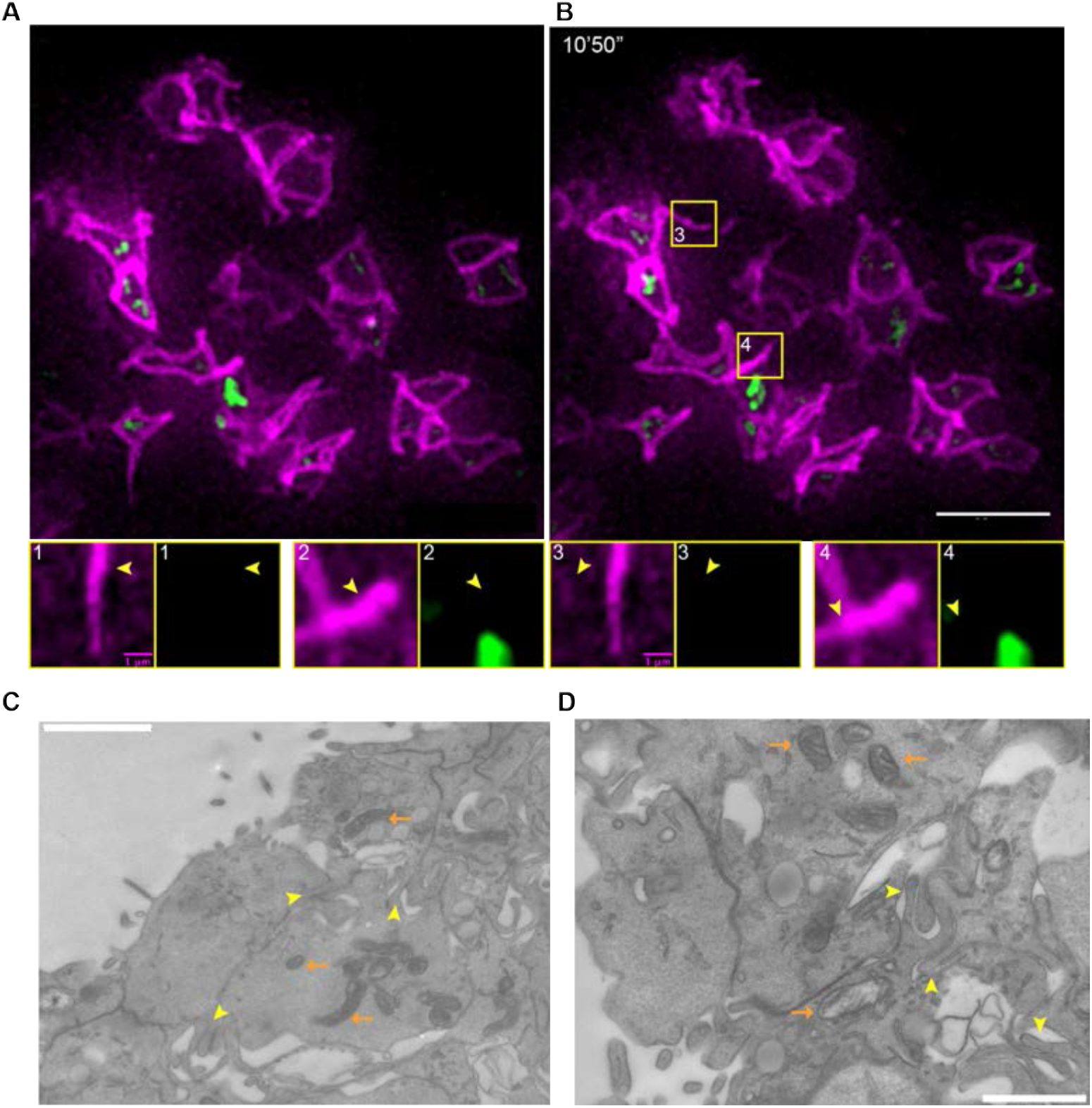
Mitochondria do not enter lateral protrusions. (A, B) Live imaging timepoints (movie 22) following mis-expression of pm-mKate2 and the mitochondrial marker mNeonGreen-TOMM20. Boxed regions of lateral protrusions are enlarged below, arrowheads indicate lateral protrusions (3 experiments, 9 explants); (C, D) enface EM sections from the apical surface of chick embryonic spinal cord at HH17-18 (sampled in 2 embryos), showing lateral protrusions (yellow arrowheads) and mitochondria (orange arrows). Scale bars (A, B) 10 μm, (C, D) 1 μm.

**Supplementary Figure 4.**
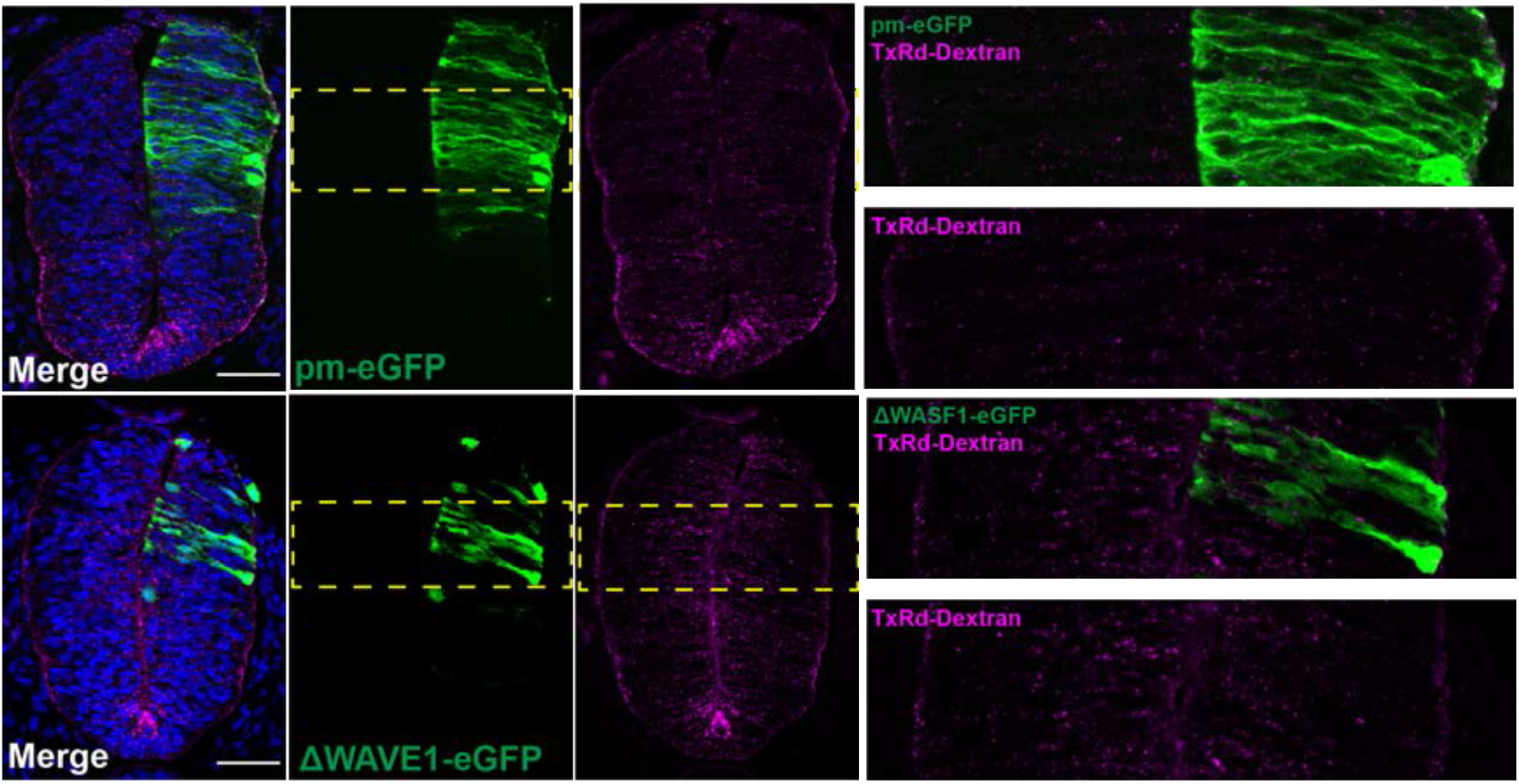
ΔWAVE1-eGFP does not compromise apical neuroepithelial integrity. Transverse sections through HH17-18 neural tube following misexpression of control pm-eGFP or ΔWASF1-eGFP and introduction of Texas Red-Dextran (70kDa) into the neural tube for 24h. No qualitative change in the levels of Texas Red-Dextran between the two conditions was observed (pm-eGFP = 4 embryos, ΔWASF1-eGFP = 4 embryos). Scale bars 50 μm

